# Functional classification of tauopathy strains reveals the role of protofilament core residues

**DOI:** 10.1101/2024.09.30.615521

**Authors:** Jaime Vaquer-Alicea, Victor A. Manon, Vaibhav Bommareddy, Peter Kunach, Ankit Gupta, Jim Monistrol, Valerie A. Perez, Hung Tri Tran, Nil Saez-Calveras, Siling Du, Sushobhna Batra, Charles L. White, Lukasz A. Joachimiak, Sarah H. Shahmoradian, Marc I. Diamond

## Abstract

Distinct tau amyloid assemblies underlie diverse tauopathies but defy rapid classification. Cell and animal experiments indicate tau functions as a prion, as different strains propagated in cells cause unique, transmissible neuropathology after inoculation. Strain amplification requires compatibility of the monomer and amyloid template. We used cryo-EM to study one cell-based YFP-tagged strain, resolving its amyloid nature. We then used sequential alanine (Ala) substitution (scan) within tau repeat domain (RD) to measure incorporation to pre-existing tau RD-YFP aggregates. This robustly discriminated strains, defining sequences critical for monomer incorporation. We then created 3R/4R or 4R WT RD (aa 246-408) biosensors. Ala scan of recombinant tau seeds with the Alzheimer’s Disease fold matched that of AD homogenate. We scanned 22 brain lysates comprising 4 tauopathies. This clustered cases by neuropathological syndrome, revealed the role of amino acids in protofilament folds, and allowed strain discrimination based on amino acid requirements for prion replication.

**Teaser:** Discrimination of tau strains based on the relative contribution of each amino acid to templated propagation of the amyloid.

## Introduction

Neurodegenerative tauopathies have diverse clinical and pathological presentations, and include Alzheimer’s disease (AD), corticobasal degeneration (CBD), chronic traumatic encephalopathy (CTE), primary age related tauopathy (PART) and progressive supranuclear palsy (PSP) (*1*). Most are sporadic, and some are caused by dominant missense mutations in the tau gene. All feature intracellular accumulation of tau protein in fibrillar amyloid assemblies. Experimental evidence now supports a prion mechanism in tau pathology progression, whereby pathogenic tau assemblies of unique structure, termed “seeds” move between cells and serve as templates for their own amplification by recruiting natively folded monomer (*2–4*). We originally observed that tau transmitted an aggregated state from the outside to the inside of a cell, and between cells (*2*). Contemporaneously, Clavaguera et al. reported that inoculation of tau preparations into mouse brain induced intracellular pathology (*5*, *6*). We subsequently determined that tau forms distinct, self-amplifying structures in clonal cell lines that created transmissible patterns of aggregate pathology in cells and mice following inoculation(*7*, *8*). Based on these experiments, we proposed that tau be considered a prion: a protein that propagates unique disease-causing conformations as “strains,” which are assemblies of unique structure that influence incubation times and patterns of transmissible pathology, and which vary across different tauopathies.(*9*). In parallel, we adapted simple cell-based assays, termed “biosensors,” to measure levels of tau seeding activity based on induced aggregation of tau fragments fused to fluorescent proteins paired for fluorescence resonance energy transfer (FRET) (*4*, *10*). Tau biosensors sensitively detect the presence of tau seeds in human, murine and recombinant protein samples, and have allowed detailed analyses of progressive seeding pathology in mice (*4*) and humans (*11–13*). Importantly, these methods are readily adaptable to high-throughput, quantitative analyses, although it has been unclear to what extent they report accurately on seed structure.

The patterns of end-stage deposition of tau filaments—locations, cell types, and pathological inclusion morphologies—have formed the “ground truth” for neuropathological classification of tauopathies (*14*). In the last several years, however, the cores of insoluble tau filaments extracted from brain have been characterized with atomic resolution by cryogenic electron microscopy (cryo-EM) (*15–20*). Unique core structures are linked to specific neuropathological syndromes, but the same structure can appear in multiple disorders, indicating that this alone cannot explain all features of individual tauopathies, and accessory factors likely play a role (*21*). For example, AD and the more slowly progressive PART feature the same protofilament fold, while the accumulation of Aβ uniquely in AD may underlie its faster progression (*22*, *23*).

The rapid and unbiased classification of tau assembly conformation is an important challenge. Cryo-EM is expensive and labor-intensive, and limited to particles that can be purified and directly visualized on EM grids. We hypothesized that it might be possible to solve this problem by exploiting the prion “species barrier” for tau, in which monomer recruitment to a specific strain requires a compatible amino acid sequence. We reasoned that specific mutations in native tau could reduce or block its incorporation into some strains, but not others. Furthermore, substitution of alanine (Ala), “scanning” through tau monomer, followed by measurement of its incorporation into pre-existing aggregates, would allow us to discern the relative contribution of each amino acid to the stability of growing tau assemblies, test the fidelity of intracellular replication of strains, and efficiently classify tauopathies *in vitro*. Consequently, we created a biosensor system that coupled systematic Ala substitution in tau monomer with measurement of its incorporation into aggregates based on FRET. This has established that prion mechanisms underlie the cellular amplification of unique tau structures, revealed the role of specific residues and in the formation of strains, and created an unbiased classification system for tauopathies.

## Results

### Design and development of an amino acid profiling system

We previously curated a library of 18 tau strains propagated in HEK293T cells that express the tau repeat domain (244-375aa, termed Tau RD) with disease-associated mutations (P301L and V337M, termed “LM”) fused to eYFP (table S1) (*7*). Each strain was previously characterized using intracellular inclusion appearance, biochemistry, limited proteolysis, and neuropathology in brain tissue following intracerebral inoculation(*7*, *8*). We first confirmed that Tau RD(LM)-YFP forms intracellular amyloids, focusing on DS13, a strain featuring inclusions with a disordered appearance (Fig. 1A) that was originally derived by seeding the Tau RD (LM)-YFP parental cell line (DS1) with clarified brain homogenate from a human CBD brain.

**Fig. 1.**
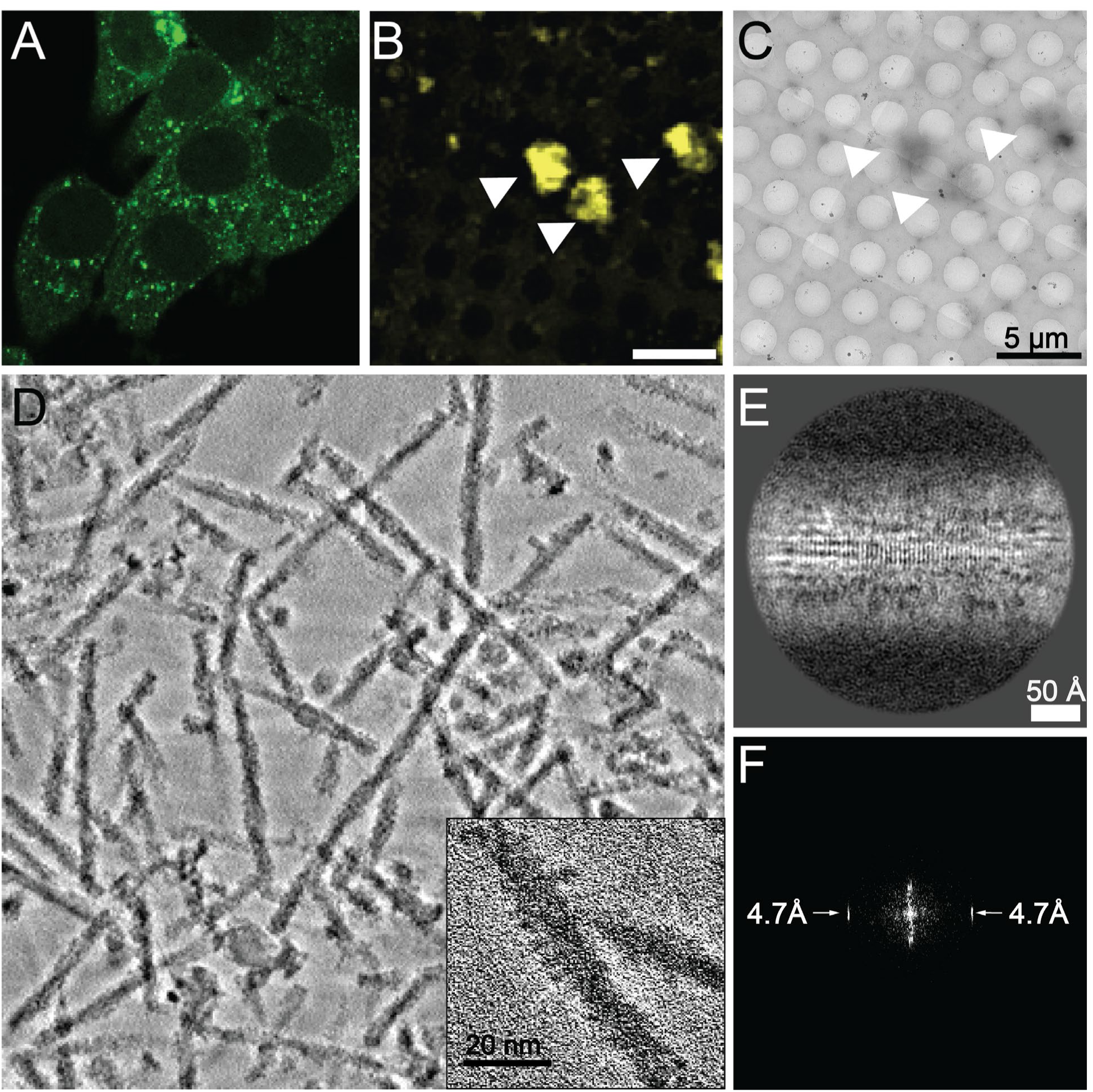
DS13 stably propagates amyloids. **(A)** Tau RD(LM)-YFP was observed to form stably propagating inclusions in DS13 cells. **(B)** Low-magnification overview of a single representative grid square imaged using cryo-fluorescence microscopy and **(C)** cryo-TEM of Tau RD(LM)-YFP inclusions purified from DS13 cells. Notably, the fluorescent signal overlapped with the high-contrast, appendages marked by arrowheads in the cryo-TEM overview. Aggregates were composed of fibrillar assemblies, as seen in **(D)** a summed projection of 152 central slices from a representative reconstructed cryo-electron tomogram, (inset) zoomed-in view of a single fibril in the tomogram and movie S1. **(E)** a 2D class average shows the cross-beta repeat of the amyloid fibril core and its corresponding FFT (F) shows a strong peak at 4.7 Å.

To confirm that the visible YFP inclusions in these cells were composed of tau amyloids, we extracted and purified insoluble protein from the DS13 cell line. Upon imaging with correlated light and electron microscopy, we observed large YFP-positive inclusions (Fig. 1B and S1A-C) which coincided with protein aggregates (Fig. 1C, S1A-C). Cryo-EM tomography indicated they were filamentous (Fig.1D). High-resolution reconstructions of the filaments using single-particle cryo-EM were hampered by difficulty with particle alignment due to the flexible moieties that decorated the surface of the fibrils (Fig. 1D, inset. S1D). We confirmed that the filaments were helical with a ∼650 Å twist (S1D). The appendages that decorated the filaments were consistent with the dimensions of eYFP. Importantly, the Fourier transform of images of individual filaments from cryo-TEM micrographs revealed a strong signal at 4.7Å, consistent with the intermolecular spacing between beta strands along the amyloid fibril axis (S1E). This was also the case for 2D-class averages of extracted filaments (Fig. 1E-F). Our data thus indicated Tau RD (LM)-YFP formed intracellular amyloid fibrils decorated with YFP.

### Measuring monomer incorporation into intracellular aggregates

We next developed a strategy for systematic mutation within tau RD to study effects on monomer incorporation into fibrils (Fig. 2). We generated an arrayed lentiviral plasmid library that encoded Tau RD (LM) fused to eCFP harboring single Ala substitutions along the length of the entire fragment (Tau RD (LM)-CFP). We validated the expression of all mutants by western blot (S2A-F). To assess the effect of each tau mutant on incorporation into aggregates, we plated each strain-containing cell line in a 96-well format and transduced it with the arrayed lentivirus library. After incubation, following incorporation of the lentiviral-encoded tau monomer into pre-existing aggregates, we harvested and analyzed the cells by flow cytometry (S3A). To rule out that changes in FRET related to protein expression of the tau-CFP variants, we confirmed similar expression of CFP fusions within the analysis gate by flow cytometry in the parental cell line DS1 (S3B). We further visually confirmed that overexpression of Tau RD (LM)-CFP variants did not induce spontaneous aggregation in the parental Tau RD (LM)-YFP cell line (DS1), which lacks aggregates, and no Ala substitution induced spontaneous aggregation of tau (S4).

**Fig. 2.**
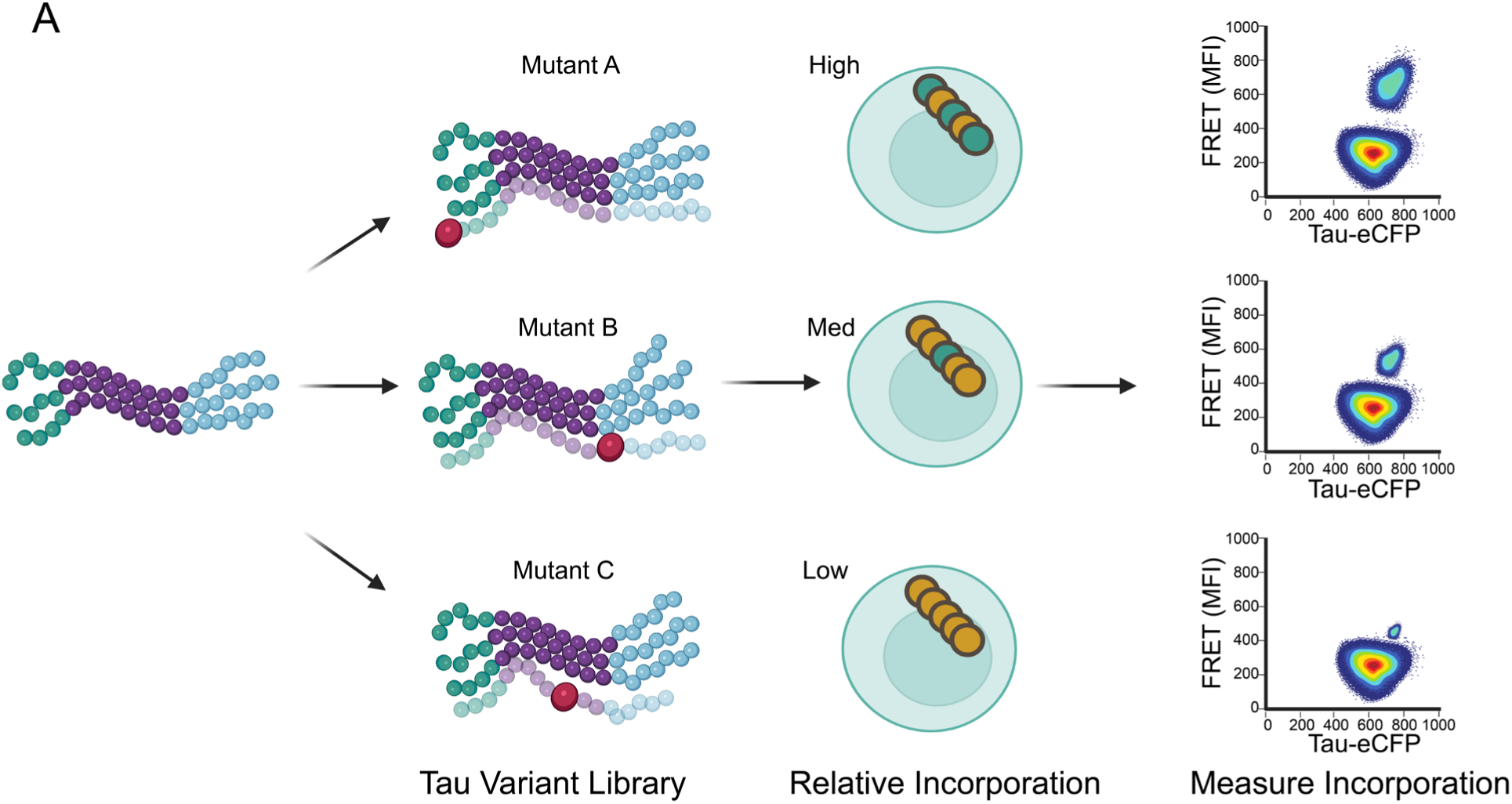
Incorporation assay overview. **(A)** The assay design relies on the expectation that mutating residues in the core of amyloid forming regions of tau will affect aggregation in a strain-dependent manner. The workflow for the incorporation assay consists of transducing cells which stably propagate tau amyloid strains with an arrayed lentiviral library to express various forms of mutant Tau RD(LM)-CFP and measuring the degree of incorporation of the mutants onto the stable inclusions propagated on Tau RD(LM)-YFP using flow cytometry.

Tau RD (LM)-CFP was invariably enriched in Tau RD (LM)-YFP inclusions when expressed in DS13 cells, which propagate a tau strain (Fig. 3A). Introduction of known amyloid-inhibiting mutations (I277P, I308P) (*24*) termed Tau RD (LM/2P)-CFP prevented incorporation (Fig.3B). We next used flow cytometry to measure the incorporation of Tau RD (LM)-CFP into growing aggregates, recording the median fluorescence intensity in the FRET channel over time (Fig. 3C). Saturation of the FRET signal past 48 hours coincided with steady-state levels of protein expression. This indicated that the incorporation of monomers into growing assemblies was likely limited by protein expression and not the aggregation process. Consequently, to limit the bias introduced by different lentiviral titers between mutants for subsequent experiments we analyzed cells with comparable levels of CFP expression (S3B). Expression of another disease associated mutant P301S in Tau RD produced similar levels of aggregation, while expression of the WT version of the fragment (lacking LM mutations present in the cell lines) had low FRET signal, presumably because of diminished incorporation (Fig. 3C).

**Fig. 3.**
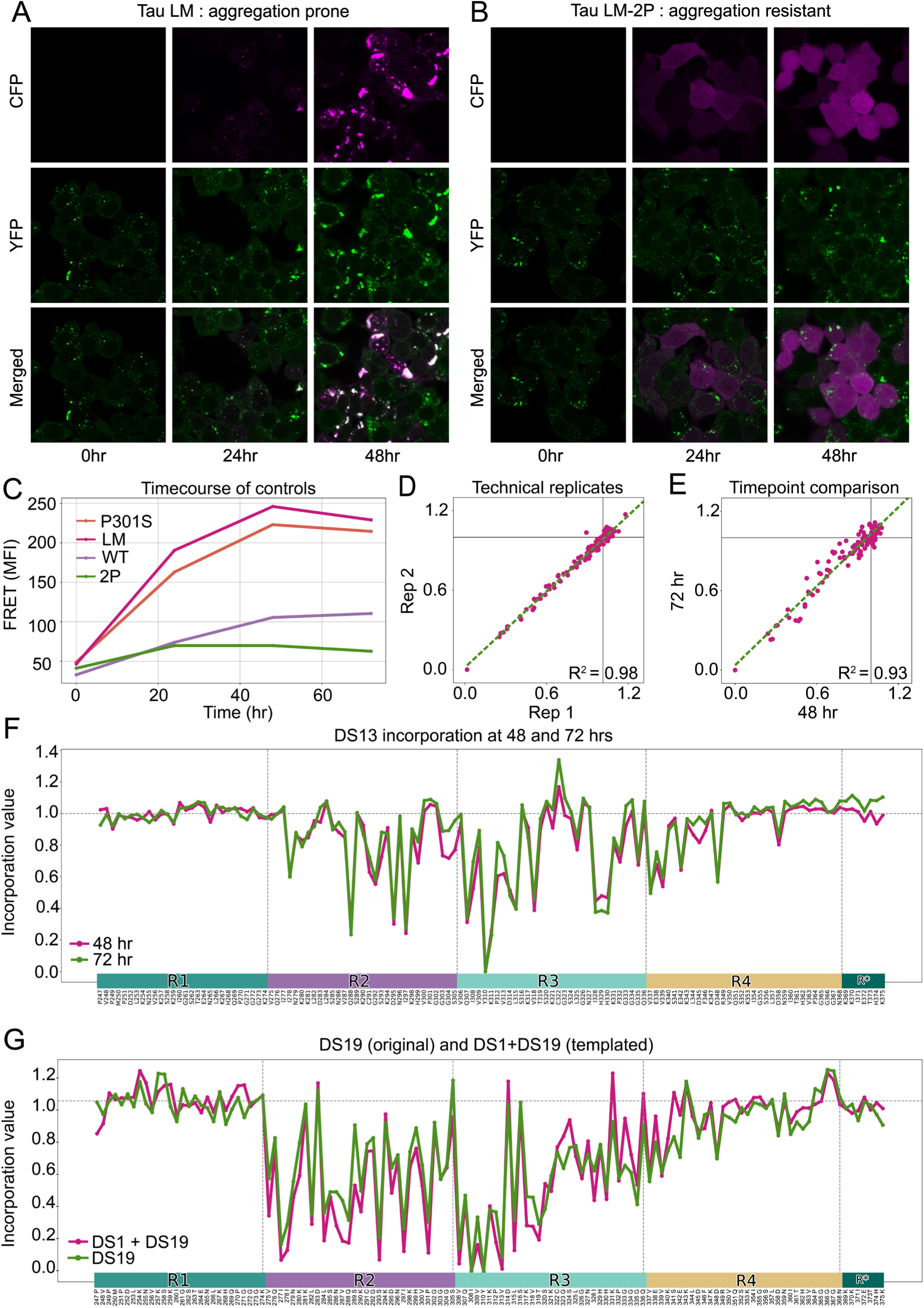
Incorporation assay reports a signature of templated aggregation. **(A)** Tau RD(LM)-CFP was detectable in DS13 cells at 24-48 hours after transduction with lentivirus. Inclusions were apparent at 24hrs and colocalized with endogenous LM-tau-YFP expressed in cells. **(B)** Cells that were transduced with Tau RD(LM/2P)-CFP lentivirus lacked inclusions only in the CFP channel. **(C)** The colocalization of CFP and YFP in inclusions within DS13 cells treated with LM-CFP lentivirus was detectable via FRET which increased over time and saturated after 48 hours. Interestingly, removal of both disease-associated mutations, rendering the WT sequence, also prevented the incorporation into DS13 aggregates. We observed a consistent, intermediate level of FRET when a different disease-associated mutation P301S was used in place of Tau RD(LM)-CFP. **(D)** Scatter plot of DS13 replicate scans performed 48 hours after transduction with lentivirus. (E) Scatter plot of incorporation assay on DS13 performed at 24 and 48 hrs. **(F)** Line plots of alanine scan performed at 48 and 72 hours of incorporation. Vertical lines demarcate repeat domain boundaries in tau. **(G)** Line plots of DS19, another strain from our library, and that of DS1 (parental line) seeded with DS19 homogenate prior to an alanine incorporation scan, were essentially identical.

### Monomer incorporation identifies predicted amyloid motifs

We then quantified incorporation of each Ala mutant into the pre-existing aggregates using flow cytometry. We observed signal intensities between Tau RD(LM) and Tau RD (LM/2P) differing by ∼5 fold (Fig. 3C), which bounded the range of signal. We next compared the levels of incorporation of each of 128 Ala substitutions into the DS13 strain. Each value for DS13 (Fig. 3D-F) represents a different Ala substitution at a given position in the tau sequence. The changes in magnitude are presented relative to the amount of Tau RD (LM)-CFP incorporated into aggregates of DS13, with values <1 representing less incorporation compared to WT Tau RD. Replicate scans indicated high reproducibility between technical replicates (R^2^=0.98) (Fig. 3D) and between measurements at two different timepoints (R^2^=0.93) (Fig. 3E-F). Ala substitutions throughout substantial portions of the first repeat (R1) and second half of the fourth repeat (R4) did not measurably impact monomer incorporation. Conversely amino acids within R2 and R3, beginning at residue 275 and ending with amino acid 358, were especially important (Fig. 3F). As expected, strong effects occurred within the hexapeptide aggregation motif ^306^VQIVYK^311^ in R3. Interestingly, Ala substitutions outside of ^306^VQIVYK^311^, particularly near the middle of the second repeat (Q288, D295 and I297) had strong effects, as did three residues further into the third repeat (I328, H329, H330), which have been identified as amyloid promoting regions (APRs) by orthogonal methods (*25*, *26*)

To confirm that changes in incorporation were not an artifact of the clonal cell lines and in fact represented structural features of the seed, we extracted filaments from another clone, DS19, and transduced DS1 cells, which do not contain inclusions, before repeating the Ala scan. We observed the same pattern of hits in DS19 and in DS1 seeded by DS19 (R^2^=0.90), indicating that information transfer was mediated by protein assembly and not the cell line (Fig. 3G). We tested selected hits by confocal microscopy and observed colocalization of CFP inclusions and YFP aggregates consistent with changes in incorporation. (S4B). Taken together, this information was consistent with stable propagation of a precise amyloid conformation from the outside to the inside of a cell.

### Synthetic tau strains differentiated by incorporation patterns

We next tested all 18 previously characterized synthetic strains to investigate their amino acid requirements for monomer incorporation. These derived from recombinant fibrils, human tauopathy cases, or a murine model of tauopathy (PS19) that were initially propagated within the DS1 line (*7*). We plated each strain in a 96-well format, prior to exposing it to the arrayed lentiviral library. We first compared strains with distinct morphological appearance as assessed by confocal microscopy (table S1). We easily distinguished DS13, a disordered strain, from DS18, a strain with ordered appearance that was originally derived from CTE homogenate (Fig. 4A). We then performed the incorporation assay on the complete library of strains and found that most strains were sensitive to different Ala substitutions that did not correlate with cellular morphological inclusion patterns (Fig. 4B, table S1). For example, incorporation into DS18 was most sensitive to substitutions in R4 while most other strains, except for DS7, mostly depended on residues in R2 and R3. Importantly, we failed to distinguish several pairs of strains previously described as different, e.g. DS3 and DS19 (Fig. 4C), both derived from recombinant fibrils. Upon closer inspection of the published data that supported their original distinction (*7*), we observed only minor differences between the DS3 and DS19 strains across all assays.

**Fig. 4.**
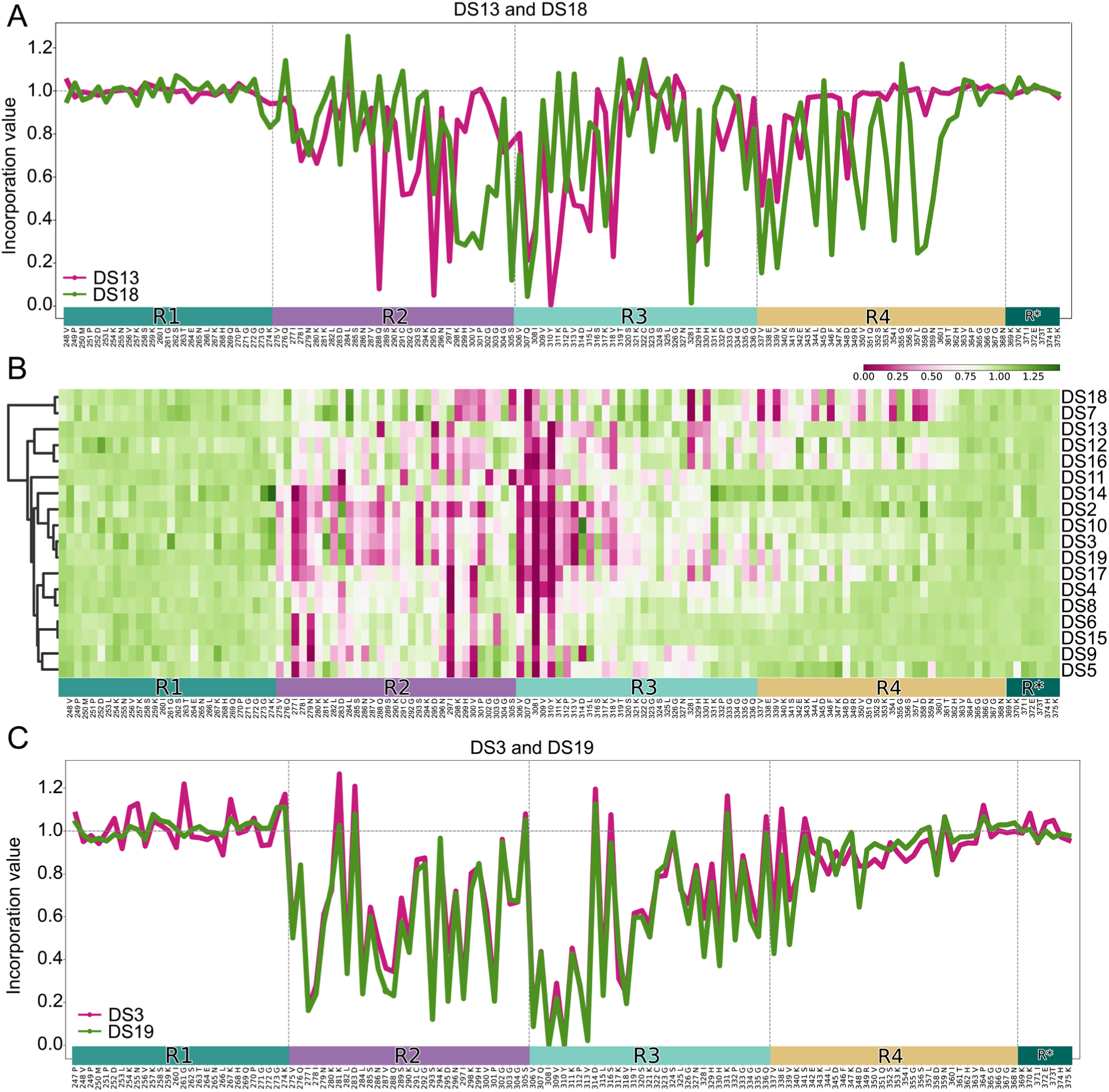
Reliable discrimination of DS strains. **(A)** Line plots of dissimilar strains, highlighting usage of diverse repeats in the tau sequence. **(B)** A heatmap of the incorporation profiles of all DS strains confirms the diversity of strains in our library. **(C)** Line plots comparing the two most similar alanine scans from our library: DS3 and DS19.

Throughout the length of the repeat domain, we found regions with patterns that involved every other residue, reminiscent of the expected alternating patterns of in-out of core side chains of an amyloid, for example, ^294^SDNIKHV^300^, in DS11 (Fig 4B). In no strain did we observe this pattern in regions predicted to be involved in turns that link beta strands, such as the four “PGGG” motifs and the four “KXGS” motifs present throughout the Tau RD (LM) fragment (S5). On average DS strains were affected by mutations starting in the second tau repeat and ending around aa 361 (S5). This contrasted with amyloid cores described in WT tau fibrils from multiple tauopathies, which have typically involved the first repeat through amino acid 386.

On average, the distribution of incorporation scores in the beginning and end of the tau RD sequence centered around 1 (S6A), which differed markedly from the other positions (S6B). While most strains used similar regions, broadly encompassing R2 and R3, the location and magnitude of hits varied widely. Interestingly, the ^306^VQIVYK^311^ sequence was critical among all strains.

Next, we used the correlations between incorporation scores to apply unbiased cluster analysis, comparing strain similarity based on the requirements for assembly (Fig. 5A). The information content of the incorporation assay clearly discriminated tau strains within cells based on template-mediated assembly. Broadly, there were two remarkably distinct sets of strains; one composed of DS7 and DS18, which stand out for their usage of R4, and a second group that encompassed the rest (Fig. 5A). Within the larger group, there were smaller sub-clusters, such as one including DS2, DS3, DS10 and DS19, that showed high intra-group correlation. The magnitude of correlations within this group exceeded those expected by chance, calculated by generating one thousand permutations of the data, randomizing the incorporation values by sample, and calculating the pairwise correlations (S6C). Hierarchical clustering revealed a strong correlation, with all major hits overlapping for DS4 and DS8; DS6 and DS15; DS7 and DS18; DS12 and DS16; DS3 and DS19. This indicated each pair is similar, if not the same structure (Fig. 4B, 5A). Based on the unbiased nature of this discrimination scheme, we conclude that probably these were originally misclassified as distinct.

**Fig. 5.**
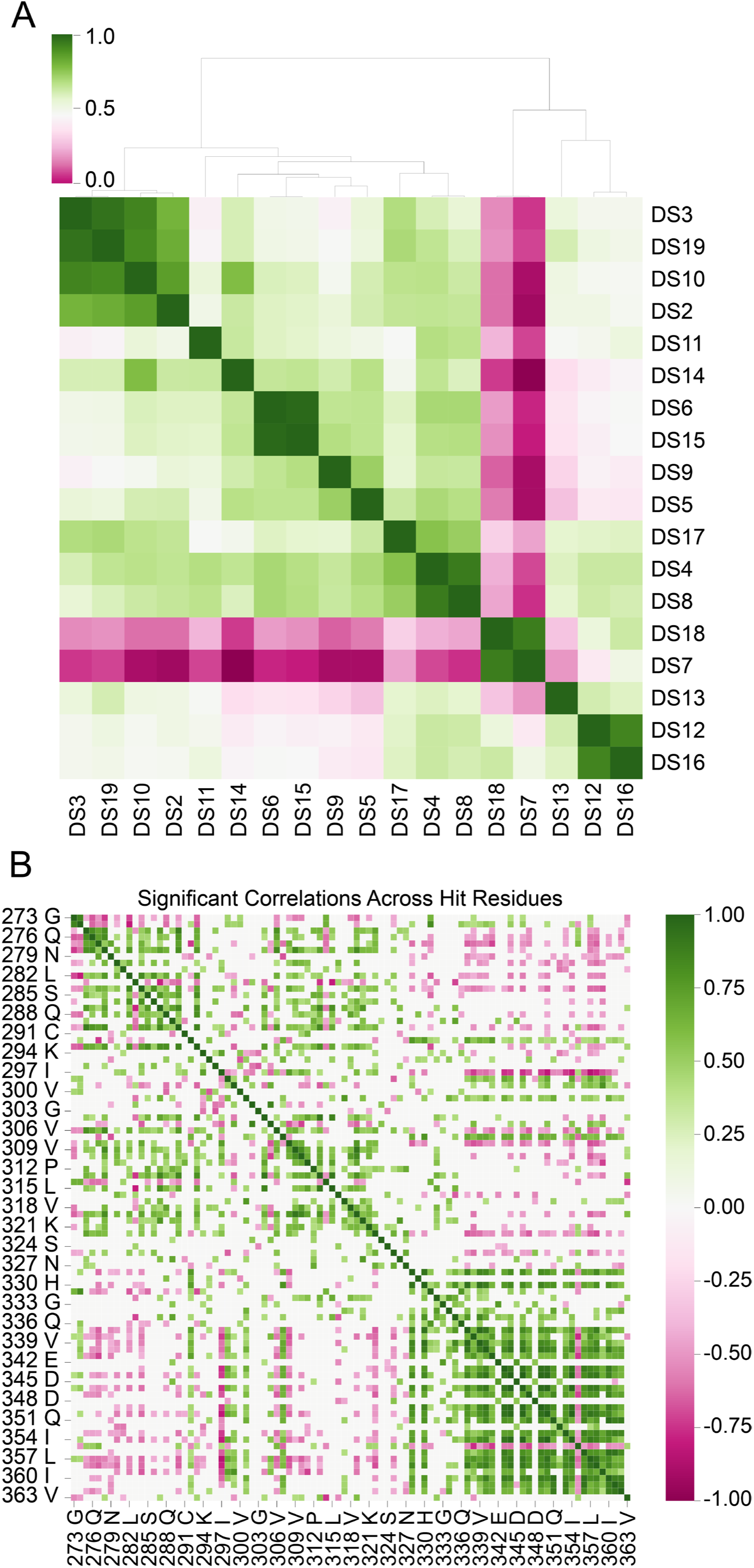
Clustering of DS strains based on incorporation assay. **(A)** Clustergram based comparison of all strains in the tau RD(LM)-YFP library. Broadly, there were two very distinct sets of strains; one which encompassed DS7 and DS18 and a second group which encompassed the rest. Within this larger group, there were smaller sub clusters that showed very high intra-group correlation, such as DS2, DS3, DS10 and DS19, highlighted in dark green. **(B)** Clustergram of the correlation coefficients for each pairwise comparison of mutated positions. This was filtered to only show correlations deemed significant (threshold at R=0.5 based on permutation test). Short range correlations were abundant especially within 274-294, 306-321 and 339-363.

Comparing residues by degree of correlation highlighted three distinct regions associated with aggregation (Fig. 5B) which coincided broadly with most of each repeat domain. We also observed examples of residues whose requirements correlated or anti-correlated with other residues, suggesting a modular architecture of strains in which one domain is used at the expense of another (Fig. 5B). For example, when residues in R2 are involved in the aggregation of strains, residues in R4 between 337 to 360 tend to not be involved. Interestingly, the junction between R2 and R3 (298 to 307) tends to correlate in usage with R4, while the adjacent position I297 is the most anticorrelated, which may suggest a regulatory role. Related observations have been made previously regarding local structural preferences being associated with amyloid polymorphisms (*26*).

### Strain-typing soluble seeds from human tauopathy

We next focused on tau seeds derived from human tauopathies. We characterized multiple extensions of the WT RD sequence to identify those that would retain efficient seeding and report on all possible known amyloid core structures. We identified an extended Tau RD sequence (aa 246-408) that we fused either to Cerulean3 or mRuby3. These were co-expressed as a novel biosensor, termed Tau RD(4Rext)-Cer/Rub. To ensure capture of 3R seeds, we created a second cell line that expressed the 3R version of the extended fragment (Δ275-305) fused to Cer along with the 4R version of this fragment fused to mRuby3 termed Tau RD(3R/4Rext)-Cer/Rub. We also generated an Ala-substituted lentiviral plasmid library with the aa 246-408 tau fragment fused to mEOS3.2 (S7A). We adapted a similar protocol to the DS strain Ala scans, except we began each assay by inducing aggregation in biosensors with tauopathy samples. Thus, we treated each cell line (4R/4R or 3R/4R) with a sample that contained tau seeds derived from either recombinant sources or human tauopathy cases, waited 48h, and replated the cells onto 96-well plates, transducing the following day with the arrayed 4R tau variants. This intermediate step was added to ensure ala variant incorporation would be measured against the same initial structure, rather than seeding directly onto each mutant, which might cause different polymorphs to form. After an additional 48-72 h to allow incorporation, we harvested cells for analysis by flow cytometry or confocal microscopy (Fig. 6A, 6B and S7B-E).

**Fig. 6.**
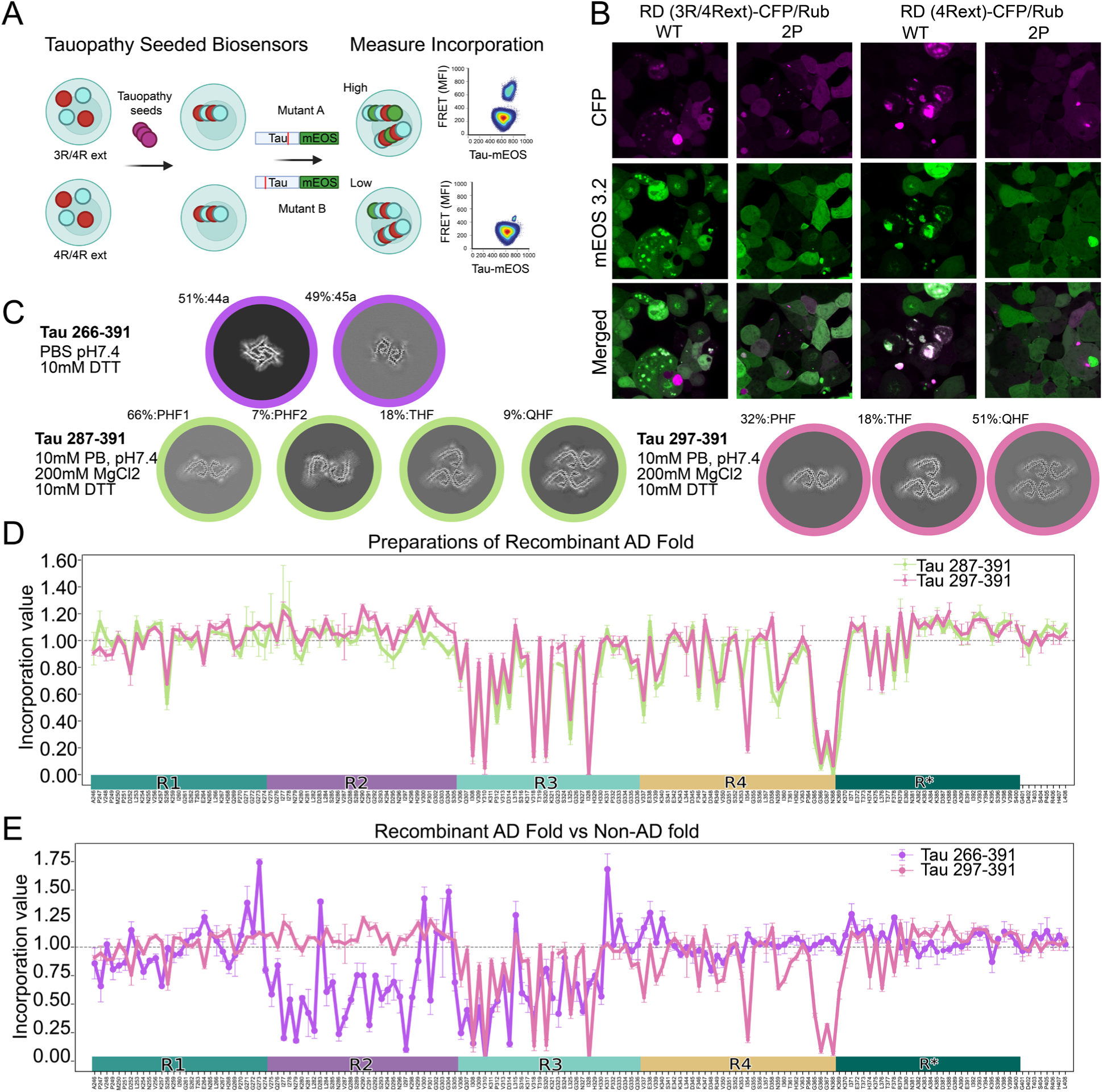
A WT-tau incorporation assay reports on amyloid cores. **(A)** We modified the incorporation assay for use with a broader source of samples. This involved seeding stable WT-tau-RD-CFP/mRuby expressing cells with test samples to induce aggregation before transduction of an arrayed WT-tau-RD-mEOS.3.2 lentiviral library, incubation, and readout via flow cytometry. **(B)** Confocal images of Tau RD(3R/4Rext)-CFP/Rub and RD(4Rext)-CFP/Rub which were seeded with AD brain homogenate and, after 48 h, transduced with either the unmodified Tau RD(4Rext/WT)-mEOS3.2 or the anti-aggregation RD(4Rext/2P)-mEOS3.2. Importantly, while the Tau RD-eCFP fusions aggregated in both cell lines, only the WT-mEOS3.2 and not the 2P-mEOS3.2 formed inclusions and colocalized with CFP. **(C)** Tau fibrils were prepared under various conditions, as in Lövestam et al., 2022 and detailed in the methods section. The dominant morphologies of these preparations were characterized by cryo-EM and cross-sections of the reconstructions are shown. One sample with tau (266–391) produced cross-sections like filaments “44a” and “45a” from Lövestam et. al (2022), these are highlighted with the purple outline and utilize the R2 domain of tau and the first half of R3, unlike the structures in the other two preparations (see also, S9D-E). The tau (287–391) fibrils, highlighted in green, and Tau (297–391) fibrils, highlighted in pink, were composed of protofilaments with the AD fold. In the first, the tau (297–391) fibrils were composed of 3 different protofilament arrangements, while 4 different arrangements of protofilaments were present in the tau (287–397) fibrils. **(D)** Tau RD(4Rext)-CFP/Rub were treated with tau (287–391) fibrils and the resulting Ala scan plotted in green. In pink, the incorporation profile of tau (297–391) fibrils which was performed in Tau RD (3R/4Rext)-CFP/Rub biosensors. **(E)** Tau RD(4Rext)-CFP/Rub was treated with tau (266–391) fibrils and the resulting Ala scan plotted in purple. Shown in pink is the incorporation profile of tau (297–391) fibrils which was performed in Tau RD (3R/4Rext)-CFP/Rub biosensors as shown in D.

### Alanine scan reports on amyloid cores

To test the precision of the incorporation assay for tau seeds, we adapted recently published protocols (*18*, *27*) to generate 3 samples of recombinant tau fibrils. We characterized these samples by cryo-EM (Figure 6C) and determined the presence in one sample of a mixture of two folds that are not disease-related, while the other two samples contained densities consistent with the “AD fold” in diverse protofilament arrangements. We modeled the atomic structure of tau into these densities, which matched the AD protofilament fold in all polymorphs of the two samples, with either 3 or 4 ultrastructural arrangements (two forms of Paired Helical Filaments: PHF1 and PHF2, Triple Helical Filaments: THF or Quadruple helical filaments: QHF) (Fig 6C and S8E-F). We then performed Ala scans using these structurally characterized samples as seeds. To test whether the specific biosensors influenced seeding, we performed an Ala scan with fibrils from the tau fragment encompassing (287–391) on 4Rext cells, while a second scan was performed in 3R/4Rext biosensors with fibrils made from the fragment encompassing 297-391. Despite differences in filament ultrastructure, scans from these samples with the AD fold were virtually identical (Fig. 6D), and easily distinguished from filaments with the non-AD conformation (Fig. 6E). This suggested the assay was likely most sensitive to the underlying fold and less to the protofilament packing, as we did not observe strong effects when mutating within the PHF interface ^332^PGGGQ^336^, nor the additional interface present in THFs and QHFs between E342 and K343. Importantly, it also suggested that, for AD samples, the Ala scan was not altered by usage of only 4R vs. 3R/4R for templating. Cryo-EM indicated that residues forming the amyloid core in the tau (266–391) sample involved residues 274-328, while the AD amyloid present in the first two preparations utilized residues 306-378. The Ala scans thus highly correlated with core structures determined by cryo-EM.

### Alanine scan correlates structures of recombinant fibrils and AD brain lysate

We further tested the predictive power of the Ala scan with disease-derived seeds. We observed virtually identical incorporation patterns for the recombinant AD-like fibrils and AD brain homogenate (Fig.7A-B). To understand the relationship between the hits in the incorporation assay and the structure of the filaments, we mapped the values onto a model of the AD fold present in recombinant PHFs, THFs and QHFs. Nearly all critical amino acids in the incorporation assay corresponded to residues within the amyloid core, and most mediated interactions between mated beta-strands, and not solvent-exposed residues (Fig.7C). This was consistent with a requirement for folding of the incoming monomer layer onto the growing protofilament as critical for addition, rather than residues involved in protofilament interfaces.

**Fig.7.**
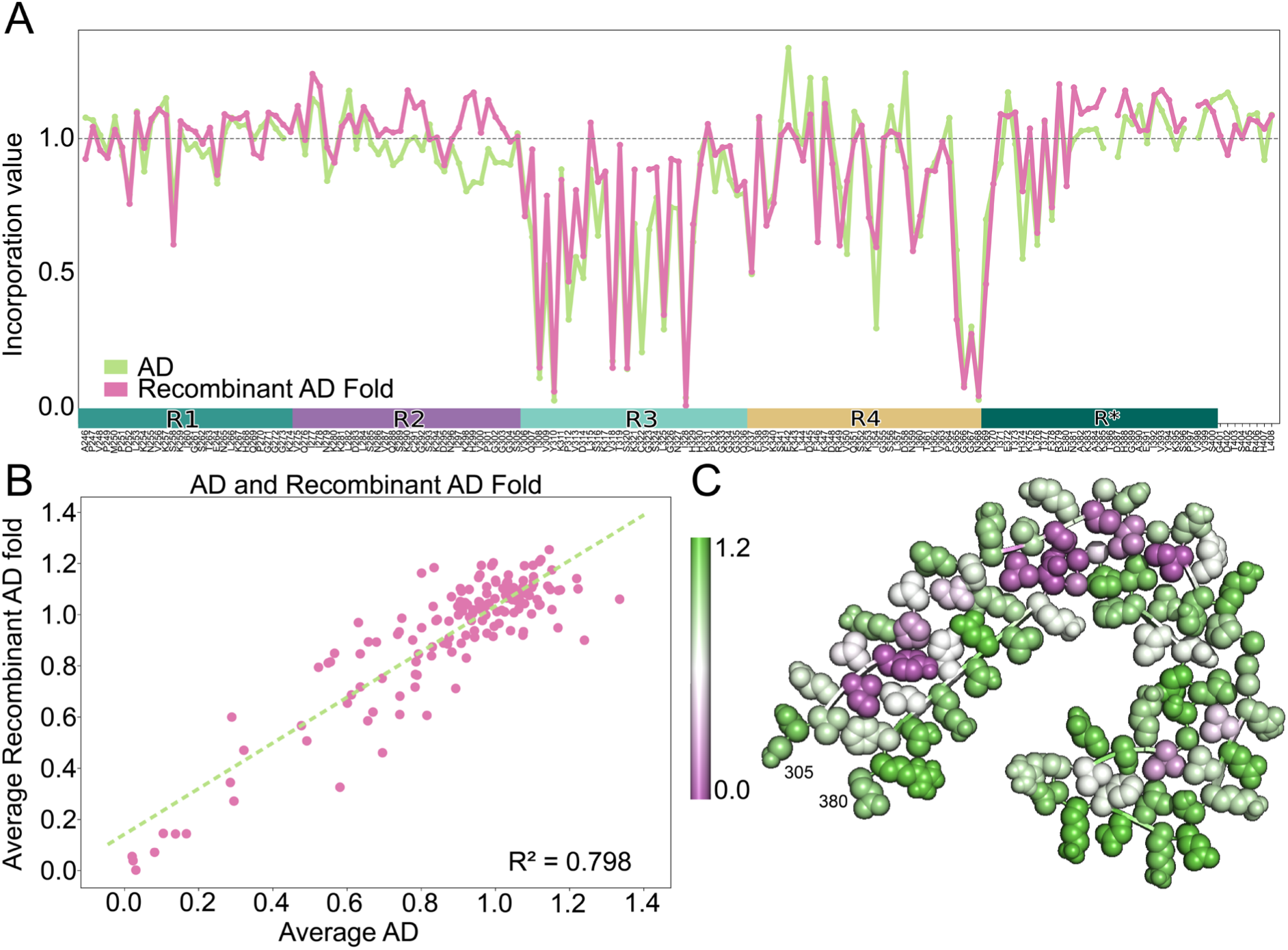
Recombinant fibrils with the AD fold mimic seeding activity of AD homogenates. **(A)** Average WT alanine scan incorporation values from samples which were composed of the AD fold reproduce patterns from AD brain homogenate. The incorporation profile of the *in vitro* fibrils correlated well **(B)** to that of AD brain homogenates. **(C)** Mapping of the incorporation assay values on the AD protofilament from PHF1 illustrates that the strongest hits were in the core of the amyloid, corresponding to residues present in the interacting faces beta sheets. Values representing no change in incorporation are in green, while positions in magenta were strongly affected by mutation.

### Incorporation assay discriminates tauopathy samples

Our results indicated that the incorporation assay reported fundamental structural features of soluble seeds. To further test this idea, we analyzed seeding from 8 AD, 5 CBD, 2 CTE and 6 PSP cases. We were restricted to cases for which seeding was strong enough to render a clear signal in the biosensors. Each incorporation assay was performed in the biosensor that resulted in the most aggregate positive cells upon seeding: CBD, PSP and CTE were read out from 4R-only biosensors, while AD was read out in mixed 3R/4R biosensors. In each case the Ala scan was performed using 4R tau incorporation into the induced pre-existing aggregates.

Seeds from different diseases required strikingly distinct components of the RD sequence. Critical residues for CBD spanned from the beginning of R2 to beyond R4, whereas those for AD began towards the end of R2 and extended past the end of R4. (Fig. 8A-D). Hits from the Ala scan correlated well with known cores observed by cryo-EM reconstructions of filaments from these disorders. Thus, as for the recombinant fibrils, the incorporation assay based on brain seeding returned accurate information about the known core structures of seeds from each condition. Notably, the pattern of hits for AD and CTE correlated well in R3 and R4, as would be expected for these similar folds, yet we observed requirement for R2 in the CTE-derived Ala scans, which had not been described as belonging to the amyloid core in disease. As has been observed in the analysis of residue energetics of amyloids (*21*, *26*), mapping incorporation values onto published cryo-EM structures of disease associated folds, we observed strong correlation of required residues within mated beta-strand pairs (Fig. 8E-H), with a notable exception again being the CTE structure which contains several solvent exposed hits within the first 10 amino acids.

**Fig.8.**
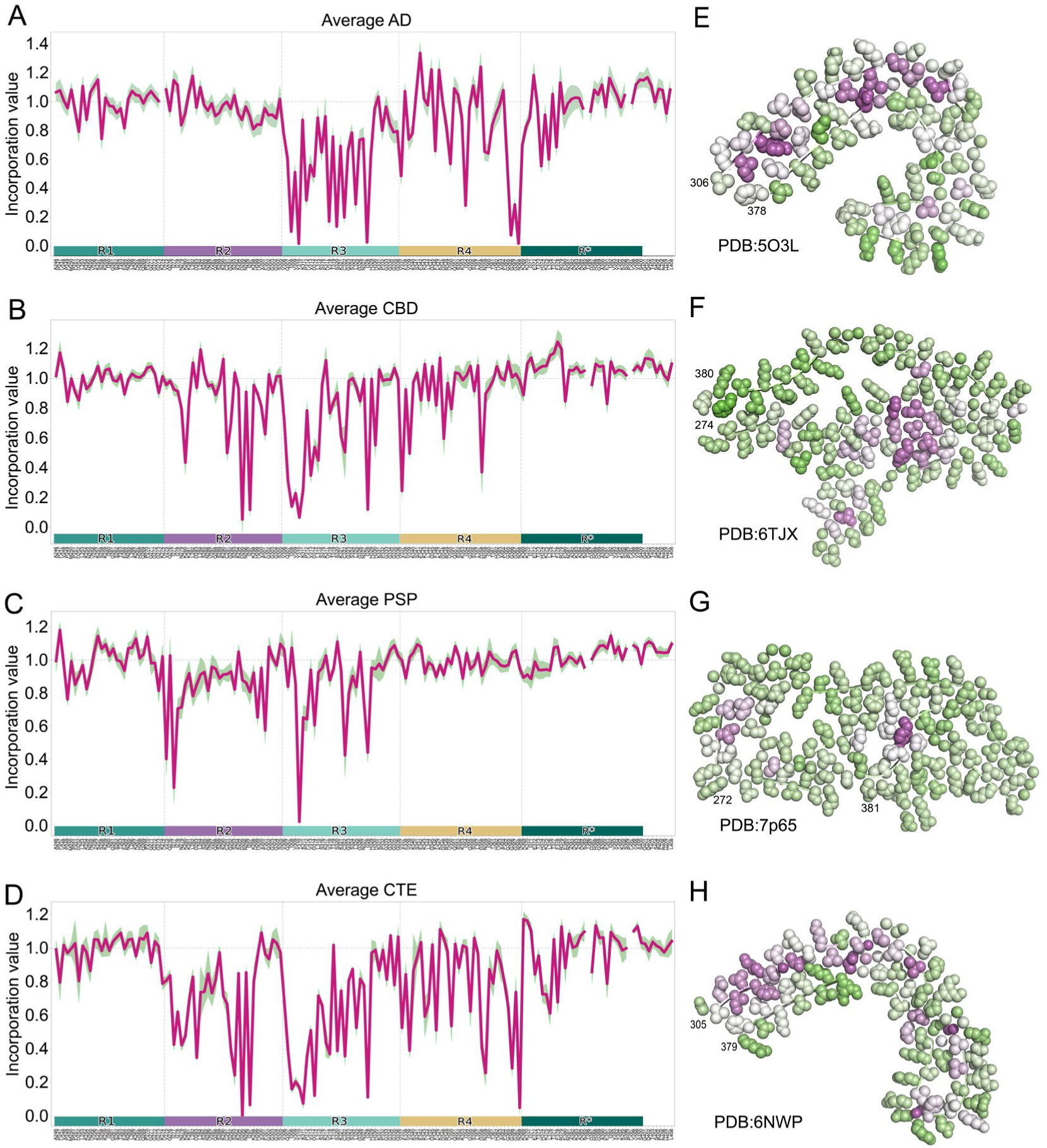
Alanine scan signature overlaps with amyloid core and identifies tauopathies. The average incorporation profile with 95% confidence interval of various tauopathy brain homogenates next to their corresponding mapped values on published models of each disease protofilament. **(A and E)** AD (n=8) mapped on PDB:5O3L, **(B and F)** CBD (n=7) mapped on PDB:6TJX, **(C and G)** PSP (n=5) mapped on PDB:7P65, and **(D and H)** CTE (n=2) mapped on PDB:6NWP. Green and Magenta represent high and low incorporation values, respectively.

Next, we compared incorporation scores across a panel of tauopathies for which the data were particularly robust: 8 AD, 7 CBD, 5 PSP, 2 CTE cases. We observed a striking similarity for groups that clustered, and this corresponded with neuropathological diagnoses (Fig. 9A). This was further supported when analyzing individual replicates for each case, where we observed all replicates clustering with their correct neuropathological diagnosis (Fig. 9B, upper). These correlations were not explained by chance given that shuffling the incorporation values of replicates across protein positions for a given neuropathological classification resulted in a median correlation coefficient of approximately 0.85 (Fig. 9B, lower). This was contrasted by the median correlation coefficient of shuffled values without regard to diagnosis of 0.50 or that of shuffled values with samples outside of the same group of about 0.30 (Fig. 9B, lower). These results indicated that the seeding activity in human samples encoded information sufficient to discriminate tauopathies based on underlying neuropathological diagnosis.

**Fig. 9.**
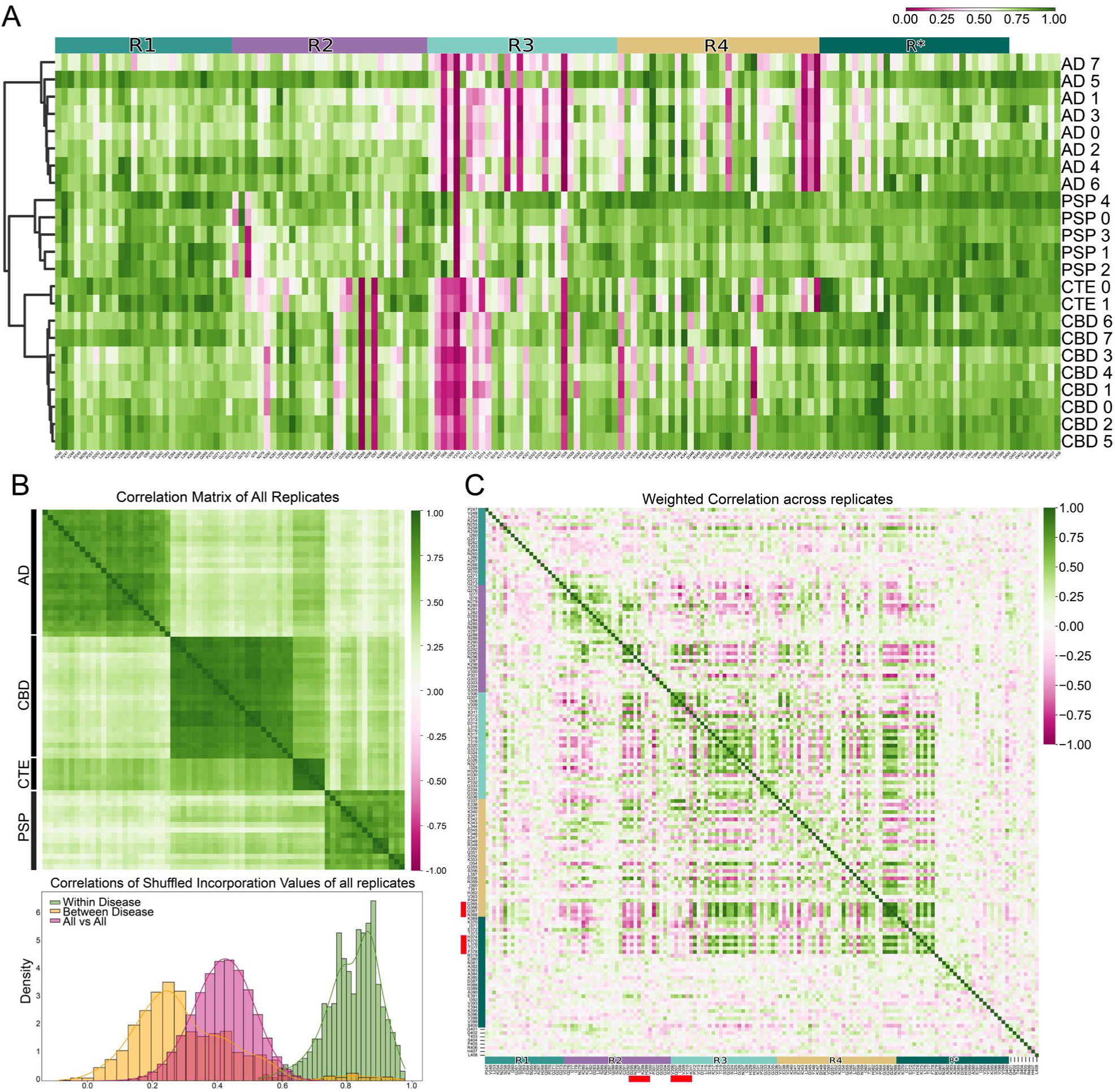
The incorporation signature clusters individuals by disease and uncovers long-range residue correlations for aggregation. **(A)** Clustered heatmap of individual cases naturally subdivides into groups that correspond to neuropathological diagnosis. **(B)** Correlation matrix of individual replicates from the twenty-three cases in A using values between 274 and 380, along with histograms of correlations between Ala scans shuffled by residue and grouped (or not) by disease. **(C)** Clustergram based comparison of the weighted correlation between residues across all treatments reveals modules involved in short and long correlations throughout the repeat domains. Highlighted in red are regions discussed in the main text.

Finally, the pairwise correlation between the incorporation values at all positions in the tau sequence highlighted regions that appear to work together to define strain identity (Fig. 9C). Briefly, most correlating hit residues lie between 271-378. Strong local correlations are evident along the diagonal in Fig.9C, dominated by signals encompassing the ^306^VQIVYK^311^ sequence, and others near ^275^VQIINK^280^ and ^295^DNIHKV^300^ in R2 and some unexpected regions involving the ^364^PGGGN^368^ turn in R4 and ^373^THKLTF^378^ just past R4. A striking pattern of long-range correlations involved D295 and I297, these negatively correlated with hits at the end of R4 and instead positively correlated with Q307, V309 and K311 in ^306^VQIVYK^311^. While the importance of this observation remains unclear, we believe this approach might help define interacting modules in tau responsible for defining strain identity and could be useful to guide structural predictions.

## Discussion

We have used seeded tau aggregation in biosensor cells to define the role of each amino acid in the incorporation of monomer into fibril assemblies. The fidelity of intracellular tau seeding allowed unbiased classification of tauopathies. We first confirmed that tau RD fusion to YFP produced intracellular amyloid fibrils with regularly spaced densities consistent with YFP. We studied a highly characterized library of individual, synthetic tau strains, formed and propagated using tau RD containing two disease-associated mutations (P301L and V337M). These served as a “ground truth” for development of an Ala scan to classify distinct assembly structures. We next produced recombinant tau fibrils whose structure we confirmed by cryo-EM—some with a protofilament fold identical to AD, and others with a fold distinct from any known tauopathy. An Ala scan using intracellular aggregates seeded by recombinant fibrils containing the AD protofilament fold was nearly identical to that performed on aggregates seeded by AD brain. This indicated high fidelity of the scan to identify seeds of defined structure. Finally, we used the Ala scan to profile multiple tauopathies, and precisely grouped them according to their neuropathological diagnosis. Thus, prion replication mechanisms in a simple biosensor were sufficient to classify the original tauopathy, based on amplification from a specific template.

### Faithful propagation of structures into cells

Prior work by the Hasegawa/Scheres/Goedert labs indicated that CBD filaments seeded into a SH-S5Y5 biosensor cell expressing 1N4R tau did not replicate the exact structure observed in disease (*28*). Intriguingly, using a variety of established biochemical measures, we had previously observed that non-physiologic tau strains stably propagated and transmitted their conformations in cultured HEK293T cells (*8*). We thus determined if the process of seeding into a cell inevitably led to an alteration in the propagated structure. Given the precision of the Ala scan, we tested the amplification of structure by scanning a tau strain, DS19, in its original line, and after it was extracted and seeded into the parental line, DS1. We observed an identical signature in the original and seeded strains. We conclude that when a particular strain is selected within a given cellular context (in this case HEK293T) it will propagate and transmit structures faithfully as would a *bona fide* prion. The reasons for this faithful propagation and replication are unknown and may depend on undefined cellular cofactors. The fidelity of this context-dependent propagation model may explain the imperfect templating results observed thus far for filaments amplified from AD and CBD on 1N4R tau. Taken together, however, our results indicate that biosensor cells contain components required to replicate certain tau strains, suggesting they could be very useful to define fundamental mechanisms of tau prion replication.

### Seed propagation fidelity is enciphered only in assembly structure

It has been unclear whether non-tau factors present in seeds, which could represent post-translational modifications, small molecules, or ligands such as carbohydrates or nucleic acid, are required for their faithful replication in cells. In at least one case, because the recombinant tau filaments comprised of the AD protofilament fold produced essentially identical Ala scans in comparison with AD brain-derived seeds, it appeared that the seed did not require an exogenous cofactor to replicate. However, this must be further tested. We created two distinct recombinant fibril preparations with identical protofilament structures vs. AD-derived filaments, albeit with distinct quaternary filament assembly patterns, and which matched those previously reported(*18*). Each of these preparations were seeded into biosensors carrying both 3R and 4R tau, or only 4R, and after inclusions formed, we measured the effect of an Ala scan on the incorporation of 4R tau. Because the scans were indistinguishable, the simplest interpretation is that each filament preparation, despite its distinct provenance (recombinant, unmodified, and without additional ligands vs. brain-derived) triggered formation of conserved folds within the cell. Indeed, this is consistent with our prior observations of faithful propagation of tau strains from mother to daughter cells, and between cells.

### Representation of human subject-derived tau assembly structure in cells

It is unknown whether replication of tau prions in cultured cells expressing tau fragments accurately represents structures found in human subjects, and indeed this has engendered some controversy (*29*, *30*). As mentioned above, a recent study probed the structure of aggregates seeded by CBD filaments, and while the induced aggregates were not identical to the starting material, aspects of the protofilament fold were quite similar, suggesting partial templating. Meanwhile, non-physiologic DS strains propagated indefinitely in cultured cells appeared to produce stable templating of aggregate structure. In the case of human tau seeds transduced into cultured cells it is unclear whether the precise protofilament fold will reproduce. Indeed, our results for CBD are most consistent with the resolved structures from seeded SH-S5Y5 cells (S9 B and C). To probe the faithfulness of templated aggregation we created recombinant tau fibrils that mimicked those found in AD. These adopted a range of filament types: paired, triple and quadruple helical filaments (PHF, THF and QHF, respectively). After confirming their structure by cryo-EM, we determined their Ala scan signatures were strikingly similar to the pattern from an AD brain homogenate. In light of the observation that SH-S5Y5 cells expressing 1N4R tau produce single protofilaments with the AD fold when seeded with AD homogenate (*28*), we cannot rule out that the HEK293T based sensors do the same, however.

Similarly, in our study of CTE, we observed hits that correlated well with regions of the core (Fig. 8H). Yet we also observed hits in R2 which is not part of the amyloid core in ex vivo filaments that were resolved by cryo-EM (Fig. 8G). In this instance, however, filaments have been shown to form *in vitro* which share the overall fold of CTE yet have a density that wraps back over the ^306^VQIVYK^311^ sequence (S9A). Consistent with this observation, a switch in isoform usage between 4R to a 3R and 4R mixture has been reported in early stages of CTE (*31*) that could account for both the observed preference for CTE seeding onto 4R biosensors and the usage of R2 that has not been observed by cryo-EM from patient samples.

The concordance of the incorporation assay with cryo-EM studies also indicated that the amyloid cores observed in filaments extracted from AD brain matched those in soluble AD extracts. A corollary is that most of the tau sequence outside the ordered core appears dispensable for templated propagation in cells. We conclude that while the HEK293T biosensor assay may not precisely replicate the structures found in humans, it clearly defines them.

### Protofilament folding defines incorporation efficiency

We were somewhat surprised to discover that monomer incorporation into induced intracellular assemblies was influenced most by residues that seemed critical to the stability of the protofilament monomer fold, and less so by generic interactions across protofilament layers. This included core residues that mediated cross-beta sheet interactions, and those involved with beta turns, consistent with previous reports (*21*, *25*, *26*). This may suggest that as a fibril grows within a cell, the incoming monomer layer may adopt a particular fold prior to, or concurrent with, its addition, and that this, and not cross-protofilament contacts, is a rate-limiting step.

### Positioning of alanine hits relates to defined filament structures

The Ala scans of defined cell-based strains and seeds derived from human brains had unanticipated power to reflect the diversity of local structures and of domains that when used in one structure were excluded from another. For example, in certain cases, we observed an alternating pattern of hits which, when correlated with structure, appeared to reflect the alternative mating of compatible residues in beta strands. In other contexts, we observed a lack of alternating hit residues around regions expected to form turns, (e.g. PGGG), potentially due to the lack of a side chain, and thus no in/out pattern. Notably, in the setting of recombinant tau fibrils which had the AD protofilament structure and a mix of quaternary structures (double, triple, and quadruple helices) we observed identical hit patterns. Finally, the Ala scan defined the core filament sequences.

### Deviation from predictive models

Not every preparation produced clearly interpretable Ala scan results. For example, one recombinant fibril preparation, tau (266–391), consisted of two distinct protofilaments mixed within the preparation. Perhaps not surprisingly, the Ala scan did not produce consistent hits within the protofilament folds as were observed with more “pure” preparations (S9D, E). In this case, the alanine scan showed a strong signal beginning in residue 274 and ending around 330. This overlapped with the amino acids identified by cryo-EM to be part of the core (44a:275-328 and 45a:274-328). While these protofilament folds used the same general sequence, packing of amino acids differed greatly between the structures. That is, some surface-exposed residues in 44a were not exposed in 45a and vice-versa. The lack of concordance in this case suggests a limitation of the assay in resolving mixtures of polymorphs. Similarly, in the case of PSP we did not observe seeding patterns that precisely matched the protofilament fold previously reported by cryo-EM(*19*), and were surprised not to observe strong hits in C-terminal residues. In this regard, it may be important that we previously identified three distinct strain composition patterns in brains from 6 human PSP subjects not studied here(*8*). Thus, when multiple structures exist in the same preparation, the Ala scan may fail to report information that allows structural classification.

### Unbiased Classification of tauopathies using soluble seeding activity

Cryo-EM has revolutionized our knowledge of filament structures within diverse tauopathies. However, cryo-EM offers limited information on the relative importance of individual amino acids outside of a well-defined core. The method reported here requires only soluble seeding activity from crude brain homogenates to classify tauopathies in a relatively unbiased fashion, with a “fingerprint” for monomer incorporation. It is of course unknown at this moment whether the Ala scan reports precisely on the conformation present in a brain-derived seed, as the assay currently relies on seeded events within cells and subsequent recruitment of monomer. But by reporting on the role of each amino acid with a relatively continuous signal range, this method offers a potentially powerful approach to classify tauopathies. Indeed, based solely on an Ala scan we easily clustered each tauopathy with similar diagnoses. The brain sample set with the greatest variance between individuals, PSP, while clearly identifiable as a discrete cluster, also may have reflected greater strain diversity within affected human subjects, as suggested by our prior work(*8*). Finally, based on these results, in the future it should be possible to design a small collection of biosensors that accurately classify most human tauopathies.

## Conclusion

This study introduces a robust, unbiased system for classification of tau strains in reductionist biosensor cells using functional genetics. This involves systematic testing of amino acid requirements for tau monomer incorporation into assemblies and has demonstrated potential for rapid and precise in vitro classification of tauopathies using little more than engineered cell lines. This work indicates that simple cell systems faithfully represent fundamental mechanisms of tau prion replication. It is very consistent with progression of tauopathy derived from mobile prion assemblies that serve as specific templates for their own replication throughout the brain, and thereby lead to defined patterns of neuropathology.

## Materials and Methods

### Cell culture

HEK293T (ATCC-CRL-1268) cells were grown in Dulbecco’s Modified Eagle’s medium (DMEM) supplemented with 10% Fetal bovine serum (HyClone), 1% Glutamax (Gibco). HEK293T cells tested negative for mycoplasma prior to making of any cell line and were grown in the absence of Penicillin/Streptomycin except for before being treated with any exogenous material, including lipofection of brain homogenate or recombinant protein.

### DS13 sarkosyl purification

Initially, the DS13 cell pellet was resuspended in a sucrose buffer composed of 0.8 M NaCl, 10% sucrose, 10 mM Tris–HCl (pH 7.4), 5 mM EDTA, and 1 mM EGTA, which has been filter-sterilized. This mixture was homogenized using a Polytron device. Following homogenization, the sample is centrifuged at 500 x g for 10 minutes, with the supernatant being preserved separately. This procedure was repeated to ensure thorough cell lysis. Subsequently, the collected supernatants were combined, and Sarkosyl added to achieve a final concentration of 2%. This mixture was then incubated with glass beads at room temperature for one hour to facilitate aggregate dissociation, employing a Nutator to ensure gentle mixing. The solution underwent ultracentrifugation at 186,000 rpm for 60 minutes to pellet the aggregates. The pellet was then resuspended in a sucrose buffer and any clumps are dispersed using a 21-gauge syringe followed by an additional ultracentrifugation step at 186,000 rpm for 60 minutes. The final pellet was resuspended in a buffer containing 20 mM Tris–HCl and 100 mM NaCl. Benzonase was added to the solution, which is then incubated at 37°C with agitation for one hour. A final ultracentrifugation at 186,000 rpm for 90 minutes was performed, and the pellet resuspended again in the purification buffer.

### Cryo-preparation of DS13

Extracted DS13 tau filaments were applied to 300-mesh copper R2/1 holey carbon grids (Quantifoil Micro Tools GmbH, Jena, Germany) that were glow-discharged for 30 s at a current of −30 mA on a PELCO easiGlowTM Glow Discharge Cleaning System. Samples were double-sided blotted with Whatman filter paper (grade 1) to remove excess liquid and then the grid was quickly plunged frozen into liquid ethane using a Vitrobot (Thermo Fisher).

### Cryo-EM data acquisition of DS13

For single-particle analysis as shown in S1D and E, 4,307 60-frame movie stacks were acquired using a Titan Krios transmission electron microscope (Thermo Fisher Scientific) operating at 300 keV and captured with a K3-Summit (Gatan) direct electron detector operating in CDS mode at a magnification of 105,000x, corresponding to a super-resolution pixel size of 0.415 Å, with a defocus range of 0.4-1.6 µm. The total dose was 60 e-/Å2. A GIF-quantum energy filter (Gatan) used in zero-loss mode with a 20 eV slit width was employed to eliminate inelastically scattered electrons.

### Cryo-EM data processing of DS13

Movies were gain-corrected, aligned, dose-weighted and averaged using MOTIONCOR2 (*32*) in RELION 4.0 (*33*). The micrographs were then CTF-estimated using CTFFIND-4.1 (*34*). Micrographs whose estimated maximum resolution was worse than 6 Å were discarded and all the resulting micrographs were manually assessed by checking the absence of ice crystals using their corresponding power spectra. Fibrils from 3,167 micrographs were then manually picked using a boxsize of 430 or 850 pixels pixels (356.9 or 705.5 Å, respectively) and 1,331,903 or 1,183,675 fragments were extracted respectively, using a 3.3X or 2X binning factor. Only 10% of the extracted fragments were used for 2D classification. In S1D.i., a regularization parameter T of 2 and a tube diameter of 150 Å were used to get a better alignment of the fibril core. In S1D.ii., a coarse alignment was obtained by setting a large tube diameter (280 Å) and a regularization parameter T of 1. The alignment of the 2D class averages was performed using the relion_helix_inimodel2D command in RELION 4.0 (*33*) with the following settings: 250 Å tube diameter and 600 Å crossover distance.

### Cryo-correlative light and electron microscopy (cryo-CLEM) imaging of DS13

Vitrified samples were imaged at -195°C utilizing the STELLARIS 5 Cryo Confocal Light Microscope from Leica Microsystems, which was equipped with a cryo-stage, Power HyD detectors, and a White Light Lasers (WLL) light source. For imaging, Z-stacks of areas displaying distinct YFP fluorescence signals were captured (excitation: 500 nm, emission: 505 nm - 732 nm) using the Cryo CLEM objective HC PL APO 50x/0.90. The imaging parameters were set as follows: z-steps of 0.2 µm, a pinhole diameter of 66.1 µm, voxel dimensions of 49 nm × 49 nm × 224 nm, pixel dwell time of 2.8125 μs, and a scan speed of 400 Hz. Additionally, images in reflection mode were captured to identify grid squares characterized by optimal ice quality, intact carbon films, and distinct landmark features crucial for correlation. The Z stacks were then exported from the LAS X software and processed in FIJI to create maximum intensity projections of the YFP signal, thereby facilitating the rapid identification of ex vivo tau-YFP aggregates on TEM grids. Correlation was subsequently performed manually during cryo-EM imaging, with navigation to the designated ROIs guided by the cryo-confocal data.

### Cryo-ET imaging and data processing of DS13

Extracted DS13 tau filament tilt series were acquired using a Titan Krios transmission electron microscope (Thermo Fisher Scientific) operating at 300 keV and captured with a K3-Summit (Gatan) direct electron detector and a Volta Phase Plate with a target defocus of −0.5 μm (*35*) . A Bioquantum post-column energy filter (Gatan) used in zero-loss mode with a 20 eV slit width was employed to eliminate inelastically scattered electrons. The microscope control software SerialEM v4.0.8 was utilized to operate the Krios and collect 25 tilt series from 60° to −60° in 2° increments using a dose-symmetric tilting scheme (*36*, *37*). Tilt series were acquired in counting mode at a magnification of 42,000x, corresponding to a final pixel size of 1.989 Å. Each tilt angle movie stack contained 4 frames collected over 0.4 s (0.1 s/frame) with a dose per tilt of 1.52 e^-^/Å^2^. Tilt series were aligned using AreTomo (*38*) and reconstructed using IMOD eTomo (*39*).

### Common plasmid backbone generation

The FUGW plasmid backbone, originally from the David Baltimore laboratory, was modified as previously described(*10*). Briefly, the sequence starting with the UBC promoter and ending with the GFP was replaced the CMV promoter and Kozak sequence (GCCGCCACCATG) followed by a BSMBI restriction enzyme cloning cassette that ends just before a linker sequence coding for a 12 amino acid linker (GSAGSAAGSGEF), a fluorescent protein (FP-which varied) and a stop codon. This backbone, referred to here as FM5-CMV-nk-cassette-linker-FP backbone was used to generate all plasmids used in this study with further modification by changing the cassette region for the coding sequences of tau and its variants, and the fluorescent proteins (FP) region for various fluorophores including mCerulean, mRuby, eCFP, eYFP or mEOS3.2.

### LM-tau alanine scan library

Site-directed mutagenesis primers were used to generate gene fragments containing alanine substitutions at each position of the human tau gene from residue 244-375 that already contained mutations at position 301 and 337 to Leucine and Methionine, respectively, generating Tau RD(LM) alanine variants. Gene fragments were inserted via Gibson cloning into the common lentiviral plasmid encoding an eCFP as a fluorescent protein. All plasmid inserts were sequence verified by Sanger sequencing.

### Lentivirus production

Lentivirus was prepared as described previously(*40*). Briefly, cells were plated at approximately 30% confluency in either 6-well, 24-well or 12-well plates, depending on the amount of virus required. Twenty-four hours later, TransIT-293 was used to transfect psPAX2 and VSVg packaging plasmids along with our common lentiviral FM5 transfer plasmid (described above) at a ratio of (3:1:1) following manufacturer’s instructions. After 48hrs of growth, cell media was collected, spun at 100x g for 5 minutes to remove cellular debris and the remaining media was aliquoted and stored at -80C for further use.

### WT-Tau (246–408)- alanine mutant library generation

Twist Biosciences synthesized gene fragments encoding the human tau sequence from residue 246 to 408 with alanine codon substitutions (GCC) at each position. These gene fragments were then cloned into our common lentiviral plasmid with the mEOS3.2 fluorescent, generating an arrayed library of plasmids. To enable future library pooling and sequencing the linker sequence between the tau gene and the fluorescent protein was doubled in size by encoding the same amino acid sequence but with different codons for each alanine mutant. This allowed for sequencing of this small region for mutant identification in a pool of sequences. All plasmids were sequence verified by Sanger sequencing. Western blots were run to using Tau A antibody (in-house antibody targeting R1 of tau).

### Cell line generation

### RD(4Rext)-Cer/Ruby

For 4R only biosensors, HEK293T cells were treated with both lentivirus containing the human Tau RD (residues 246–408), C-terminally fused to a monomeric Cerulean3 (Cer), and monomeric Ruby fluorescent protein (mRuby3) to constitutively express the WT tauRD-Cer protein and WT tauRD-mRuby. Forty-eight hours following treatment, single cells were sorted into 96-well plates after the expression of the protein was confirmed by epifluorescence microscopy. The cell lines were tested for seeding with CBD brain homogenate and the most sensitive clone was expanded, frozen and stored for future use in incorporation experiments.

### RD(3R/4Rext)-Cer/Rub

For mixed tau isoform cells, the 3R version of the tau fragment (246–408) was C-terminally fused to a monomeric cerulean (Cer), and 4R to monomeric Ruby fluorescent protein (mRuby3). Forty-eight hours following treatment, single cells were sorted into 96-well plates after the expression of the protein was confirmed. The cell lines were tested for seeding with AD brain homogenate and the most sensitive clone was expanded, frozen and stored for future use in incorporation experiments.

### Protein expression and purification

Sequence-verified plasmids were transformed into E.coli BL21(DE3) cells from Agilent, for protein expression using standard methods and plated on 10cm^2^ agar plates. After overnight incubation at 37°C, a single colony was picked and inoculated into 500mL of lysogeny broth auto-induction media in a beveled flask and grown while shaking at 300rpm for 8hrs at 37°C in a NewBrunswick Innova 43/43R shaker. This was followed by expression for 16 hours at 24°C. Cells were harvested by centrifugation at 4,000xG for 20minutes at 4°C. Cell pellets were then frozen and stored at -80°C until use. Frozen pellets were thawed on ice and incubated with at 10 ml/g of pellet with washing buffer (WB): 50 mM MES at pH 6.0, 10 mM EDTA, 10 mM DTT, supplemented with 0.1 mM PMSF and cOmplete EDTA-free protease cocktail inhibitors. Lysis was performed by passing resuspended bacterial cells through a PandaPlus Homogenizer running at 15,000 PSI until homogenized. The sample was kept on ice throughout this procedure to prevent prolonged overheating. Lysed cells were centrifuged at 15,000x g for 40 minutes and filtered using a 0.2μm cut-off filter. The sample was then injected into 3 sequential HiTrap SP HP 5ml (15ml total) columns (GE healthcare). For cation exchange, the column was washed with 10 column volumes of WB and eluted using a gradient of WB containing 0–1 M NaCl. Fractions of 1mL were collected and analyzed by SDS-PAGE followed by staining with SimplyBlue stain. Protein containing fractions were pooled and concentrated in a 15ml Amicon centrifugal filter unit. While concentrating, samples were buffer exchanged into 10mM Phosphate Buffer pH 7.2, with 10mM DTT and loaded on a 10/300 Superdex 75 size exclusion chromatography column. Fractions were again collected and analyzed by SDS-PAGE followed by SimplyBlue Staining. When necessary, both IEX and SEC were repeated to obtain highly pure protein. Pure fractions were concentrated with a 3kDa cut-off filter and used immediately. We verified the mass of each protein purification. The intact mass of samples was verified by liquid chromatography coupled to mass spectrometry. This analysis was performed by the UTSW Proteomics core using a Sciex X500B QTOF mass spectrometer coupled to an Agilent 1290 Infinity II LC with a POROS R1 column for protein separation.

### Tau fragment fibrillization

We performed *in vitro* fibrilization reactions for all 3 fragments based similar conditions to those tested by Lövestam et. al(*18*). In all cases, 10mg/mL of protein was incubated with appropriate buffer and salts (detailed below) and incubated in a total volume of 150uL per well of a 96-well black polystyrene plate. The plate was sealed with UV-transparent film and incubated with shaking (200rpm) on an Omega plate reader (BMG Labtech) at 37°C for 48hrs. Replicate reactions were included with the addition of 5uM Thioflavin T dye to monitor amyloid formation.

Fibril preparations conditions:

1: Tau (266–391) in Phosphate Buffer pH 7.4 with 10mM DTT and 200mM NaCl

2: Tau (287–391) in Phosphate Buffer pH 7.4 with 10mM DTT and 200mM MgCl2

3: Tau (297–391) in Phosphate Buffer pH 7.4 with 10mM DTT and 200mM MgCl2

### Cryo-EM sample preparation for recombinant fibrils

Samples with a good distribution of fibrils as evidenced by uranyl acetate negative stain TEM were prepared for cryo-EM data collection. For cryo-EM, a volume of 3.5 µL of the sample was applied to a glow-discharged holey carbon grid (Quantifoil Cu R1.2/1.3, 300 mesh) and rapidly frozen in liquid ethane using an FEI Vitrobot Mark IV (Thermo Fischer Scientific). The grids were then loaded onto a 300 kV Titan Krios microscope for imaging. Movies were acquired using a Gatan K3 summit detector in CDS mode. The super resolution pixel size was set at 0.415Å. To eliminate inelastically scattered electrons, a GIF quantum energy filter (Gatan) with a slit width of 20 eV was employed. Specifics for number of frames and electron dose along with other details can be found in table S3.

### Datasets collected and acquisition parameters for recombinant fibrils

All datasets were acquired at the UTSW Cryo-electron microscopy facility on a Titan Krios microscope. Datasets and parameters are detailed in table S3.

### Helical Reconstruction for recombinant fibrils

The collected movies were binned by two, motion corrected, and dose weighted using the motion correction implementation provided by RELION 4.0 (*41*). Aligned and dose-weighted micrographs were then used to estimate the contrast transfer function (CTF) using CTFFIND-4.1 (*34*). Initially, filaments were manually picked and extracted using a box size of approximately 440 Å with an inter-box distance of 14.25 Å (three subunits), which was then downscaled by a factor of four. A first round of reference-free 2D classification was performed using a mask diameter of 440 Å and a tube diameter of 200 Å, resulting in 2D-class averages where polymorphs could be distinguished. The highest resolution averages for each polymorph were selected, and particles were re-extracted using a box size of 280 Å without rescaling. The best 2D-class averages were chosen combined and used for inimodel generation. For de novo 3D initial model generation, the relion_helix_inimodel2d program (*42*) was employed with a cross-over distance that varied depending on the polymorph. A first 3D auto-refinement was conducted using the initial model as a reference. Subsequently, a second 3D auto-refinement was performed using a soft-edged solvent-flattened mask (lowpass filtered, 15 Å) generated from the output of the first refinement. Finally, the appropriate symmetry was applied depending on the polymorph and a search range for helical rise and twist was set to accurately refine the half-maps. Bayesian polishing was performed on the particles, and the polished particles were used for a fourth round of 3D auto-refinement. Three rounds of CTF-refinement were subsequently conducted. Following this, a 3D classification without further image alignment was performed to eliminate segments that contributed to suboptimal 3D reconstructions. For further refinement, a single class containing was selected. These particles underwent another set of auto-refinement, particle polishing, and three rounds of CTF refinements to generate a final reconstruction. Masking and post-processing techniques were applied, resulting in maps described in table S3, using an FSC criterion of 0.143. The reconstruction incorporated the appropriate symmetry, twist, and rise using the relion_helix_toolbox program(*42*), with a central Z length of 30%. The post-processed map obtained from the above steps was then utilized for model building and refinement. Additional details can be found in table S3.

### Model Building and Refinement for recombinant fibrils

A model consisting of three rungs was initially generated using the deposited model 5o3l (*17*). An individual rung was then extracted from the model using Chimera (*43*) and subjected to real-space refinement with ISolde (*44*). Model validation was performed using Phenix (*45*). Additional information and details can be found in table S3.

### Human brain sample collection

AD, CBD and PSP human brain tissue was obtained through the University of Texas Southwestern Medical Center Neuropathology Brain Bank with Institutional Review Board (IRB) approval. Select AD and PSP and CDB cases were obtained from the Alzheimer’s Disease Research Center (ADRC) from Washington University in St. Louis and CTE cases were obtained from the CTE Center at Boston University. A summary of cases is detailed in table S2.

### Tissue preparation

Human sample preparation from fresh frozen tissue was performed as follows. Frozen brain material was suspended in tris-buffered saline (TBS) containing cOmplete mini protease inhibitor tablet (Roche) at a concentration of 10% w/vol. Homogenates were made by probe homogenization with and Power Gen 125 tissue homogenizer (Fischer Scientific) and homogenates were then centrifuged at 21,000 x g for 20 min. The supernatant was retained as the total soluble protein lysate. Protein concentration was measured with the Pierce 660 assay (Pierce). Fractions were aliquoted and stored at -80°C prior to immunoprecipitation and seeding experiments.

### Transduction of biosensor cells with recombinant and brain derived seeds

For treatment with recombinant filaments, biosensors cells were plated at a density of 20,000 cells per well of a 96 well plate. Twenty-four hours later, they were transduced with a complex of 2uL recombinant filaments, 0.75uL of Lipofectamine 2000 (Invitrogen) and 17.25uL of Opti-MEM for a final treatment volume of 20uL per well. For treatment with brain homogenates, we optimized the treatment conditions to maximize the number of cells with aggregates while minimizing toxicity from treatment. For this we treated with between 5ug and 20ug of 10% brain homogenate and reduced the volume of Opti-MEM correspondingly to maintain a 20uL treatment volume per well.

### Confocal imaging of DS strains and seeded biosensors

For LM strains, 20,000 cells per well grown in phenol-red free media and were plated in PDL-coated 96-well glass-like bottom plates (Cell-Vis) and on day one treated with sufficient virus to infect >80% of cells (∼15uL). After 48hrs, cells were stained with Draq5 and imaged in an InCell 6000 high-throughput confocal microscope under DAPI and FITC channels. TIFF images were imported into ImageJ, image contrast was adjusted to 100-2500 for the FITC channel and 100-7500 for the DAPI channel, images were cropped to one fourth of their original size and saved as Jpeg for display. The contrast adjustment results in some pixel saturation of which coincides with aggregates but allows for visualization of diffuse tau that is otherwise difficult to visualize due to the very strong signal of inclusions.

For Tau RD (WT) Biosensors, 20,000 cells per well grown in phenol-red free media were plated in PDL-coated 96-well glass-like bottom plates (Cell-Vis) and on day one treated with sufficient virus to infect >80% of cells (∼15uL). Each subsequent day a fresh lentivirus aliquot was used to treat a new well. Confocal images of the Cer, CFP, YFP and mEOS3.2 channels were obtained using a Zeiss inverted LSM 780.

### Incorporation assay

For LM incorporation assays were performed by plating 15,000 cells per well in 96-well plates and treating with sufficient lentivirus to achieve a transduction efficiency of >80% (3-15uL of lentivirus containing media). Replicate runs were conducted in parallel. Cells were analyzed after 48hrs. For time course experiments, cells were plated on day one and treated with fresh aliquots of lentivirus each day for the next 5 days and all cells were harvested together for analysis. For WT incorporation assays, biosensors were plated at 20,000 cells per well in 1.5 96-well plates and treated with brain homogenates or recombinant seeds as described above. After 48hrs, cells were replated 1:6 into 6 new 96 well plates and treated with sufficient lentivirus to achieve >80% transduction efficiency (10-30uL of media containing lentivirus). For both incorporation assays, cells were harvested by 0.25% trypsin digestion for 5 min at 37C, quenched with cell media, transferred to 96-well U-bottom plates, and centrifuged for 5 min at 200×*g*. The cells were then fixed in PBS with 2% paraformaldehyde for 10 min, before a final centrifugation step and resuspension in 150 µL of PBS.

### Flow Cytometry

Flow cytometry A BD LSRFortessa was used to perform FRET flow cytometry. To measure CFP or Cer, and FRET, cells were excited with the 405 nm laser, and fluorescence was captured with a 488/50 nm and 525/50 nm filter. To measure YFP and mEOS3.2, cells were excited with a 488nm laser and fluorescence was captured with a 525/50 nm filter. To measure mRuby, cells were excited with a 561nm laser and fluorescence was detected using a 610nm filter with a 20nm bandpass.

### Alanine scan data analysis

### Gating strategy for LM alanine scan

Events consistent with size and distribution of HEK293T cells were selected for analysis. These were gated for homogenous side-scattering and forward scattering. Then, a slice through the center of the YFP (GFP channel) population was selected to measure cells with similar levels of YFP expression. Lastly, the median fluorescence intensity of the FRET channel through a slice of high CFP intensity (PacBlue channel) was recorded.

After completing a full scan, values were plate normalized by dividing by the average of either the first 20 wells of the plate (for mutations 244-342 in LM) or by the last 10 residues in tau (for mutations 265-375 and controls). These values were selected because no hits were consistently observed in these areas for any strain. Next, the replicates were averaged, and standard deviations calculated. Values were considered hits if they were below 2 standard deviations from the values of mutations used as normalization values.

### Gating strategy for WT-tau alanine scans

Events consistent with size and distribution of HEK293 cells were selected for analysis. These were gated for homogenous side-scattering and forward scattering. FRET between the Cerulean and Ruby, measured in the PacBlue and Qdot605 channels, was used to gate for cells with aggregates. Within this population, a narrow gate of cells positive for FITC was selected within which FRET between Cerulean and mEOS3.2 was plotted. Within this population, a narrow slice of bright cells were selected and the median fluorescence intensity for the AmCyan channel was recorded. All gates were kept constant in the analysis of the incorporation of all tau variants in a sample. These raw values we used for downstream analysis.

### Data processing for WT scans

Raw FRET values between the Cerulean and mEOS were normalized by plate to reduce batch variation. Briefly, a constant number was added to every value in a run to prevent negative values which occur spuriously in some samples with low FRET signal. Given most mutated residues in beginning and end of the tau sequence are not important for aggregation, we used these, along with the lowest value per plate (normally corresponding to either I308A or 2P) for normalization. Thus, from each value, we subtracted the minimum value on the plate, followed by dividing by the average of background subtracted values from the first 20 residues (for the first plate of a scan-which contains the first 96 positions) or by dividing by the average of the last 10 background subtracted values (in the case of the second plate). Finally, the average of technical replicates was used for downstream analysis.

Following our original manuscript submission we found the initial values for N296A as an inhibitory mutation to be mistaken due to technical error. Thus, an exception to this analysis pipeline was made for N296A for the 8 AD cases which we found to be mistaken. For these, we repeated a smaller version of the alanine scan with 7 other tau variants (I277A, D296A, S320A, I354A, N368A, WT and 2P) that were used in combination with the originally collected data to linearly interpolate the N296A values for each of the replicates of the AD cases. N296A has no effect on monomer recruitment into aggregates seeded by AD homogenates.

### Exclusion criteria

For the LM Ala scans 204 out ∼11,039 (1.8%) data points and for the WT Ala scans 232 out of 12,551 (1.8%) values were excluded post hoc due to low or no event numbers. These were usually due to clogs during flow cytometer runs which reduced the number of events detected below 300 events or mistakes during sample handling. Missing values have been highlighted in the attached excel file. For nearly all cases in which there was a missing value, there is a replicate that Ala scan on the same sample that contains the corresponding measurement.

## Acknowledgments

We thank the Cryo-EM core facility (CEMF) and the Structural Biology Lab (SBL) at UT Southwestern Medical Center, which are partially supported by grant RP220582 from the Cancer Prevention & Research Institute of Texas, for support with Cryo-EM studies. We especially thank, Zhe (James) Chen and Yang Li from SBL for assistance with data collection. Further we would like to thank Daniel Stoddard, Raymond Welch, and Jose Martinez from CEMF for their assistance with data collection. For cell sorting and flow cytometry instrumentation support, we acknowledge the Moody Foundation Flow Cytometry Facility. We also acknowledge the assistance of the UT Southwestern Electron Microscopy Core, funded by the NIH grants 1S10OD021685-01A1 and 1S10OD020103 01. We especially thank Phoebe Doss for assistance with negative staining and TEM support. We thank Nikos Louros for comments on the manuscript.

## Author contributions

Conceptualization: VAM, JVA, MID

Methodology: VAM, JVA, VB,

Investigation: VAM, JVA, VB, PK, AG, JM, VAP, HTT, SD, SB CLW

Visualization: JVA, SHS, JM, MID

Supervision: LAJ, SHS, MID

Writing—original draft: JVA, MID

Writing—review & editing: JVA, MID, LAJ, CLW, SHS, VAP

## Data and materials availability

All data are available without any restriction upon request to the authors or is available in the main text or the supplementary materials. Density maps for single particle cryoEM datasets has been deposited at the Electron Microscopy Data Bank (EMDB) under accession codes EMD-46908, EMD-46911, EMD-46915 and EMD-46909 for PHF1, THF, QHF and PHF2, respectively, in the condition with Tau (287–391) with Phosphate Buffer, pH 7.4, 10mM DTT and 200mM MgCl2. Maps associated with PHF, THF and QHF of Tau (297–391) with Phosphate Buffer, pH 7.4 with 10mM DTT and 200mM MgCl2 the following accession codes, EMD-46870, EMD-46885 and EMD-46904. Maps associated with Tau (266–391) with Phosphate Buffer, pH 7.4, 10mM DTT and 200mM NaCl can be found under EMD-47092 and EMD-46937.

## References

1. G. G. Kovacs, Molecular pathology of neurodegenerative diseases: principles and practice. J. Clin. Pathol. 72, 725–735 (2019).

2. B. Frost, R. L. Jacks, M. I. Diamond, Propagation of tau misfolding from the outside to the inside of a cell. J. Biol. Chem. 284, 12845–12852 (2009).

3. H. Mirbaha, B. B. Holmes, D. W. Sanders, J. Bieschke, M. I. Diamond, Tau Trimers Are the Minimal Propagation Unit Spontaneously Internalized to Seed Intracellular Aggregation. J. Biol. Chem. 290, 14893–14903 (2015).

4. B. B. Holmes, J. L. Furman, T. E. Mahan, T. R. Yamasaki, H. Mirbaha, W. C. Eades, L. Belaygorod, N. J. Cairns, D. M. Holtzman, M. I. Diamond, Proteopathic tau seeding predicts tauopathy in vivo. Proc. Natl. Acad. Sci. 111 (2014).

5. F. Clavaguera, T. Bolmont, R. A. Crowther, D. Abramowski, S. Frank, A. Probst, G. Fraser, A. K. Stalder, M. Beibel, M. Staufenbiel, M. Jucker, M. Goedert, M. Tolnay, Transmission and spreading of tauopathy in transgenic mouse brain. Nat. Cell Biol. 11, 909–913 (2009).

6. F. Clavaguera, H. Akatsu, G. Fraser, R. A. Crowther, S. Frank, J. Hench, A. Probst, D. T. Winkler, J. Reichwald, M. Staufenbiel, B. Ghetti, M. Goedert, M. Tolnay, Brain homogenates from human tauopathies induce tau inclusions in mouse brain. Proc. Natl. Acad. Sci. 110, 9535–9540 (2013).

7. S. K. Kaufman, D. W. Sanders, T. L. Thomas, A. J. Ruchinskas, J. Vaquer-Alicea, A. M. Sharma, T. M. Miller, M. I. Diamond, Tau Prion Strains Dictate Patterns of Cell Pathology, Progression Rate, and Regional Vulnerability In Vivo. Neuron 92, 796–812 (2016).

8. A. D. W. Sanders, S. K. Kaufman, S. L. DeVos, A. M. Sharma, H. Mirbaha, A. Li, S. J. Barker, A. C. Foley, J. R. Thorpe, L. C. Serpell, T. M. Miller, L. T. Grinberg, W. W. Seeley, M. I. Diamond, Distinct tau prion strains propagate in cells and mice and define different tauopathies. Neuron 82, 1271–1288 (2014).

8. J. Collinge, Mammalian prions and their wider relevance in neurodegenerative diseases. Nature 539, 217–226 (2016).

9. B. D. Hitt, J. Vaquer-Alicea, V. A. Manon, J. D. Beaver, O. M. Kashmer, J. N. Garcia, M. I. Diamond, Ultrasensitive tau biosensor cells detect no seeding in Alzheimer’s disease CSF. Acta Neuropathol. Commun. 9, 99 (2021).

10. J. L. Furman, J. Vaquer-Alicea, C. L. White, N. J. Cairns, P. T. Nelson, M. I. Diamond, Widespread tau seeding activity at early Braak stages. Acta Neuropathol. (Berl.) 133, 91– 100 (2017).

11. B. E. Stopschinski, K. Del Tredici, S.-J. Estill-Terpack, E. Ghebremdehin, F. F. Yu, H. Braak, M. I. Diamond, Anatomic survey of seeding in Alzheimer’s disease brains reveals unexpected patterns. Acta Neuropathol. Commun. 9, 164 (2021).

12. S. K. Kaufman, K. Del Tredici, T. L. Thomas, H. Braak, M. I. Diamond, Tau seeding activity begins in the transentorhinal/entorhinal regions and anticipates phospho-tau pathology in Alzheimer’s disease and PART. Acta Neuropathol. (Berl.) 136, 57–67 (2018).

13. G. G. Kovacs, Invited review: Neuropathology of tauopathies: principles and practice. Neuropathol. Appl. Neurobiol. 41, 3–23 (2015).

14. B. Falcon, W. Zhang, A. G. Murzin, G. Murshudov, H. J. Garringer, R. Vidal, R. A. Crowther, B. Ghetti, S. H. W. Scheres, M. Goedert, Structures of filaments from Pick’s disease reveal a novel tau protein fold. Nature 561, 137–140 (2018).

15. B. Falcon, W. Zhang, M. Schweighauser, A. G. Murzin, R. Vidal, H. J. Garringer, B. Ghetti, S. H. W. Scheres, M. Goedert, Tau filaments from multiple cases of sporadic and inherited Alzheimer’s disease adopt a common fold. Acta Neuropathol. (Berl.) 136, 699–708 (2018).

16. A. W. P. Fitzpatrick, B. Falcon, S. He, A. G. Murzin, G. Murshudov, H. J. Garringer, R. A. Crowther, B. Ghetti, M. Goedert, S. H. W. Scheres, Cryo-EM structures of tau filaments from Alzheimer’s disease. Nature 547, 185–190 (2017).

17. S. Lövestam, F. A. Koh, B. van Knippenberg, A. Kotecha, A. G. Murzin, M. Goedert, S. H. Scheres, Assembly of recombinant tau into filaments identical to those of Alzheimer’s disease and chronic traumatic encephalopathy. eLife 11, e76494 (2022).

18. Y. Shi, W. Zhang, Y. Yang, A. G. Murzin, B. Falcon, A. Kotecha, M. van Beers, A. Tarutani, F. Kametani, H. J. Garringer, R. Vidal, G. I. Hallinan, T. Lashley, Y. Saito, S. Murayama, M. Yoshida, H. Tanaka, A. Kakita, T. Ikeuchi, A. C. Robinson, D. M. A. Mann, G. G. Kovacs, T. Revesz, B. Ghetti, M. Hasegawa, M. Goedert, S. H. W. Scheres, Structure-based classification of tauopathies. Nature 598, 359–363 (2021).

19. W. Zhang, B. Falcon, A. G. Murzin, J. Fan, R. A. Crowther, M. Goedert, S. H. Scheres, Heparin-induced tau filaments are polymorphic and differ from those in Alzheimer’s and Pick’s diseases. eLife 8, e43584 (2019).

20. N. Louros, J. Schymkowitz, F. Rousseau, Mechanisms and pathology of protein misfolding and aggregation. Nat. Rev. Mol. Cell Biol. 24, 912–933 (2023).

21. W. J. Lee, J. A. Brown, H. R. Kim, R. La Joie, H. Cho, C. H. Lyoo, G. D. Rabinovici, J.-K. Seong, W. W. Seeley, Regional Aβ-tau interactions promote onset and acceleration of Alzheimer’s disease tau spreading. Neuron 110, 1932–1943.e5 (2022).

22. R. E. Bennett, S. L. DeVos, S. Dujardin, B. Corjuc, R. Gor, J. Gonzalez, A. D. Roe, M. P. Frosch, R. Pitstick, G. A. Carlson, B. T. Hyman, Enhanced Tau Aggregation in the Presence of Amyloid β. Am. J. Pathol. 187, 1601–1612 (2017).

23. M. Von Bergen, S. Barghorn, L. Li, A. Marx, J. Biernat, E.-M. Mandelkow, E. Mandelkow, Mutations of Tau Protein in Frontotemporal Dementia Promote Aggregation of Paired Helical Filaments by Enhancing Local β-Structure. J. Biol. Chem. 276, 48165–48174 (2001).

24. R. van der Kant, N. Louros, J. Schymkowitz, F. Rousseau, Thermodynamic analysis of amyloid fibril structures reveals a common framework for stability in amyloid polymorphs. Struct. Lond. Engl. 1993 30, 1178–1189.e3 (2022).

25. N. Louros, M. Wilkinson, G. Tsaka, M. Ramakers, C. Morelli, T. Garcia, R. Gallardo, S. D’Haeyer, V. Goossens, D. Audenaert, D. R. Thal, I. R. Mackenzie, R. Rademakers, N. A. Ranson, S. E. Radford, F. Rousseau, J. Schymkowitz, Local structural preferences in shaping tau amyloid polymorphism. Nat. Commun. 15, 1028 (2024).

26. S. Lövestam, S. H. W. Scheres, High-throughput cryo-EM structure determination of amyloids. Faraday Discuss. 240, 243–260 (2022).

27. A. Tarutani, S. Lövestam, X. Zhang, A. Kotecha, A. C. Robinson, D. M. A. Mann, Y. Saito, S. Murayama, T. Tomita, M. Goedert, S. H. W. Scheres, M. Hasegawa, Cryo-EM structures of tau filaments from SH-SY5Y cells seeded with brain extracts from cases of Alzheimer’s disease and corticobasal degeneration. FEBS Open Bio 13, 1394–1404 (2023).

28. S. Kaniyappan, K. Tepper, J. Biernat, R. R. Chandupatla, S. Hübschmann, S. Irsen, S. Bicher, C. Klatt, E.-M. Mandelkow, E. Mandelkow, FRET-based Tau seeding assay does not represent prion-like templated assembly of Tau filaments. Mol. Neurodegener. 15, 39 (2020).

29. A. Mudher, M. Colin, S. Dujardin, M. Medina, I. Dewachter, S. M. Alavi Naini, E.-M. Mandelkow, E. Mandelkow, L. Buée, M. Goedert, J.-P. Brion, What is the evidence that tau pathology spreads through prion-like propagation? Acta Neuropathol. Commun. 5, 99 (2017).

30. J. D. Cherry, C. D. Esnault, Z. H. Baucom, Y. Tripodis, B. R. Huber, V. E. Alvarez, T. D. Stein, D. W. Dickson, A. C. McKee, Tau isoforms are differentially expressed across the hippocampus in chronic traumatic encephalopathy and Alzheimer’s disease. Acta Neuropathol. Commun. 9, 86 (2021).

31. S. Q. Zheng, E. Palovcak, J.-P. Armache, K. A. Verba, Y. Cheng, D. A. Agard, MotionCor2: anisotropic correction of beam-induced motion for improved cryo-electron microscopy. Nat. Methods 14, 331–332 (2017).

32. D. Kimanius, L. Dong, G. Sharov, T. Nakane, S. H. W. Scheres, New tools for automated cryo-EM single-particle analysis in RELION-4.0. Biochem. J. 478, 4169–4185 (2021).

33. A. Rohou, N. Grigorieff, CTFFIND4: Fast and accurate defocus estimation from electron micrographs. J. Struct. Biol. 192, 216–221 (2015).

34. R. Danev, D. Tegunov, W. Baumeister, Using the Volta phase plate with defocus for cryo-EM single particle analysis. eLife 6, e23006 (2017).

35. D. N. Mastronarde, Automated electron microscope tomography using robust prediction of specimen movements. J. Struct. Biol. 152, 36–51 (2005).

36. W. J. H. Hagen, W. Wan, J. A. G. Briggs, Implementation of a cryo-electron tomography tilt-scheme optimized for high resolution subtomogram averaging. J. Struct. Biol. 197, 191– 198 (2017).

37. S. Zheng, G. Wolff, G. Greenan, Z. Chen, F. G. A. Faas, M. Bárcena, A. J. Koster, Y. Cheng, D. A. Agard, AreTomo: An integrated software package for automated marker-free, motion-corrected cryo-electron tomographic alignment and reconstruction. J. Struct. Biol. X 6, 100068 (2022).

38. D. N. Mastronarde, S. R. Held, Automated tilt series alignment and tomographic reconstruction in IMOD. J. Struct. Biol. 197, 102–113 (2017).

39. C. Lois, E. J. Hong, S. Pease, E. J. Brown, D. Baltimore, Germline Transmission and Tissue-Specific Expression of Transgenes Delivered by Lentiviral Vectors. Science 295, 868–872 (2002).

40. S. He, S. H. W. Scheres, Helical reconstruction in RELION. J. Struct. Biol. 198, 163–176 (2017).

41. S. H. W. Scheres, Amyloid structure determination in *RELION* -3.1. Acta Crystallogr. Sect. Struct. Biol. 76, 94–101 (2020).

42. E. F. Pettersen, T. D. Goddard, C. C. Huang, G. S. Couch, D. M. Greenblatt, E. C. Meng, T. E. Ferrin, UCSF Chimera—A visualization system for exploratory research and analysis. J. Comput. Chem. 25, 1605–1612 (2004).

43. T. I. Croll, *ISOLDE* : a physically realistic environment for model building into low-resolution electron-density maps. Acta Crystallogr. Sect. Struct. Biol. 74, 519–530 (2018).

44. P. D. Adams, P. V. Afonine, G. Bunkóczi, V. B. Chen, I. W. Davis, N. Echols, J. J. Headd, L.-W. Hung, G. J. Kapral, R. W. Grosse-Kunstleve, A. J. McCoy, N. W. Moriarty, R. Oeffner, R. J. Read, D. C. Richardson, J. S. Richardson, T. C. Terwilliger, P. H. Zwart, *PHENIX* : a comprehensive Python-based system for macromolecular structure solution. Acta Crystallogr. D Biol. Crystallogr. 66, 213–221 (2010).

