## Supplementary Materials for "Functional classification of tauopathy strains reveals the role of protofilament core residues"

Vaquer-Alicea *et al.*

**This PDF file includes:**

Supplementary Text

Figs. S1 to S9

Tables S1 to S3

**Other Supplementary Materials for this manuscript include the following:**

Movies S1

Data S1

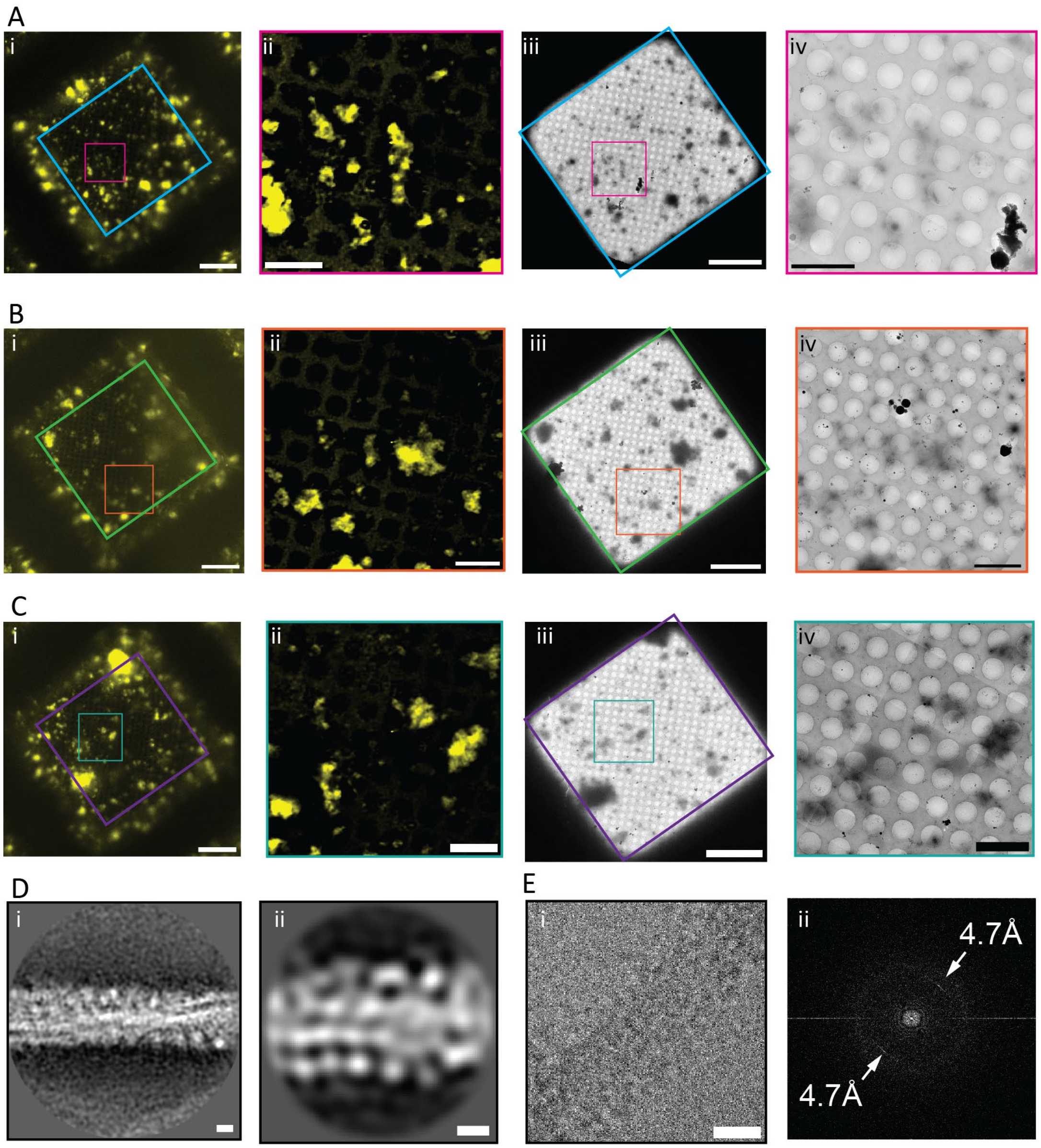
A
i
ii
iii
iv
B
i
ii
iii
iv
C
i
ii
iii
iv
D
i
ii
E
i
ii
4.7Å
4.7Å

**Sup. Fig. 1. Correlative cryo-light and electron microscopy of *ex vivo* tau-YFP aggregates.** (A-C) Additional examples showing the correlation between the YFP fluorescence (i, ii) and the electron density (iii, iv) of the tau-YFP filamentous assemblies. Images obtained by cryo-confocal microscopy (i,ii) and cryo-electron microscopy (iii, iv), respectively. Scale bars: 20 µm (i, iii), 5 µm (ii, iv). (D) 2D class averages of YFP-tagged tau amyloid fibrils. When the particles were finely aligned (i), the fibril twist (~650 Å crossover) was clearly recognizable whereas the YFP decoration was dim. When the particles were coarsely aligned, with less accuracy with respect to one another (ii), bright dots densely decorated the tau amyloid fibril and could be distinguished from each other in the 2D class average. (E) A representative micrograph with a filament along the diagonal (i) and its corresponding Fast Fourier Transform (FFT, ii). A peak at 4.7 Å, which is a feature of amyloid structure, is highlighted in the FFT.

A Tau RD-LM (247A:270A)

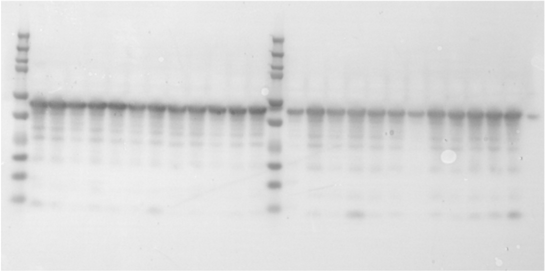

B Tau RD-LM (271A:294A)

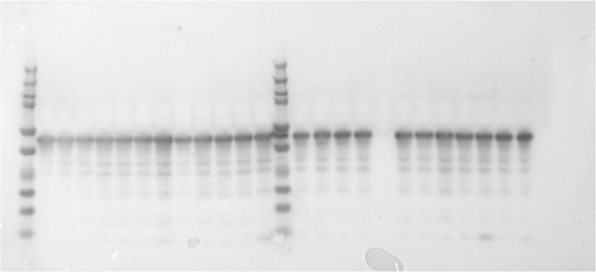

C Tau RD-LM (295A:318A)

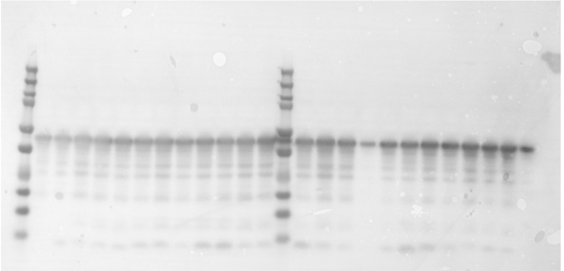

D Tau RD-LM (319A:342A)

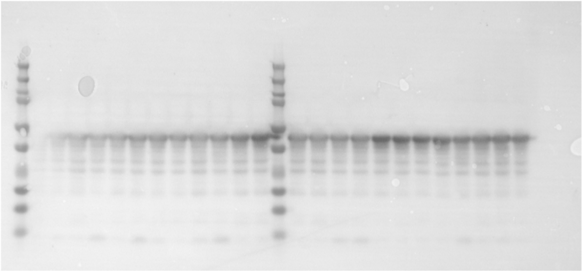

E Tau RD-LM (343A:366A)

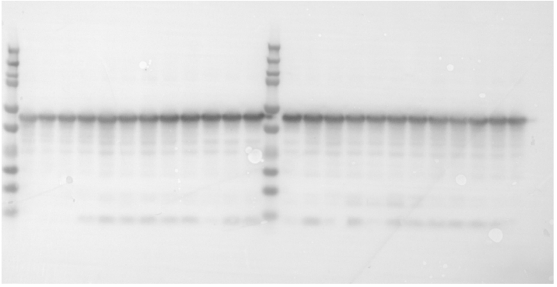

F Tau RD-LM (367A:375A:controls)

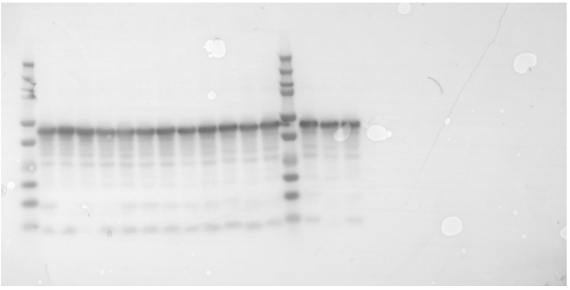

**Sup. Fig. 2 Validation of intact protein expression of all mutants and control.** Western blots of 10µg of clarified cell lysate from HEK293T cells expressing each alanine variant demonstrated robust expression of fusion protein. Lanes were loaded with each variant in consecutive order. Tau protein was detected using an in-house rabbit polyclonal antibody, named Tau A, which targets R1.

A

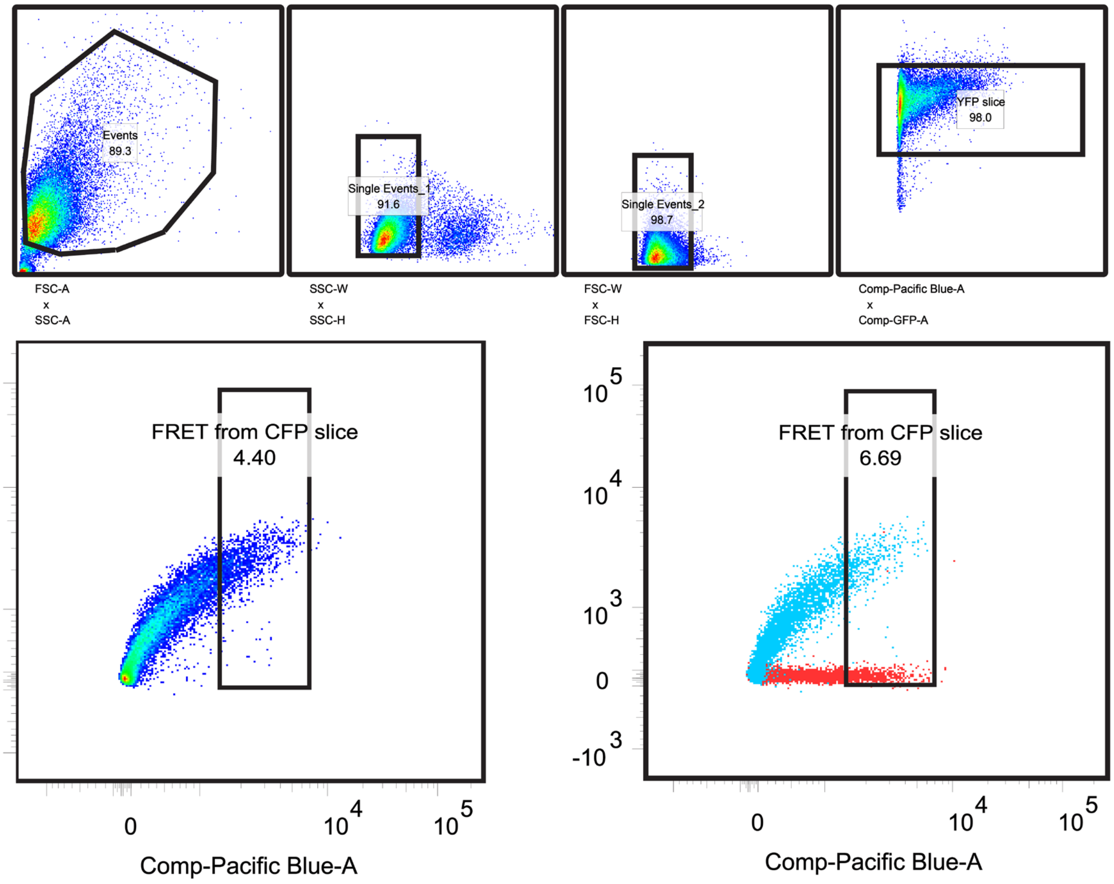
Events
89.3
Single Events_1
91.6
Single Events_2
98.7
YFP slice
98.0
FSC-A
x
SSC-A
SSC-W
x
SSC-H
FSC-W
x
FSC-H
Comp-Pacific Blue-A
x
Comp-GFP-A
FRET from CFP slice
4.40
FRET from CFP slice
6.69
Comp-AmCyan-A
Comp-Pacific Blue-A
Comp-Pacific Blue-A

B

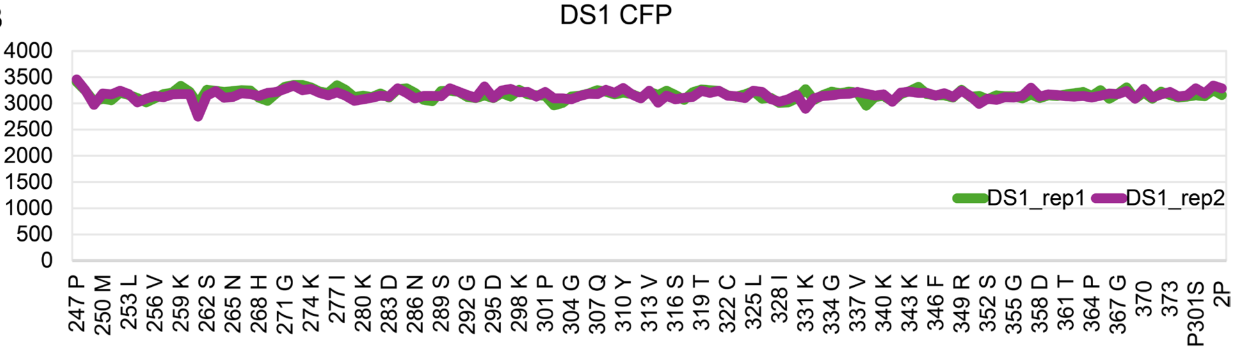
DS1 CFP
4000
3500
3000
2500
2000
1500
1000
500
0
DS1_rep1
DS1_rep2
247 P
250 M
253 L
256 V
259 K
262 S
265 N
268 H
271 G
274 K
277 I
280 K
283 D
286 N
289 S
292 G
295 D
298 K
301 P
304 G
307 Q
310 Y
313 V
316 S
319 T
322 C
325 L
328 I
331 K
334 G
337 V
340 K
343 K
346 F
349 R
352 S
355 G
358 D
361 T
364 P
367 G
370
373
P301S
2P

**Sup. Fig. 3. Gating strategy for LM Ala scan.** Events consistent with size and distribution of HEK293 cells were selected for analysis. These were gated for homogenous side-scattering and forward scattering. Then a slice through the center of the YFP (GFP channel) population was selected to measure cells with similar levels of YFP expression. Last the median fluorescence intensity of the FRET channel (AmCyan) through a slice of high CFP intensity (Pacific Blue channel) was recorded. In blue, an example of the gate used to quantify the MFI of a positive control Tau RD(LM)-CFP. An anti-aggregation negative control, Tau RD(LM/2P)-CFP, is shown in red to demonstrate the degree of signal to noise. (B) The median fluorescent intensity of CFP measured in the Pacific Blue channel of the gated population showed relatively homogeneous levels of expression of all variants.

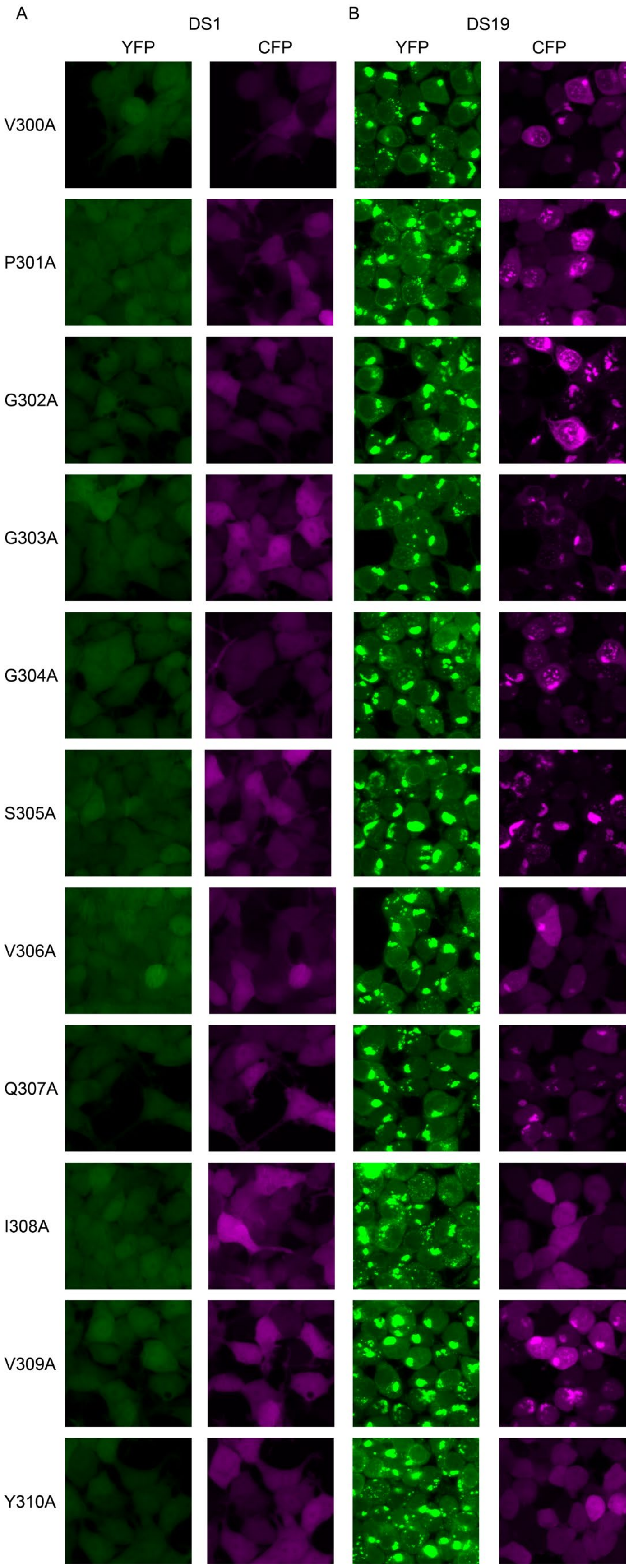
A
DS1
YFP
CFP
B
DS19
YFP
CFP
V300A
P301A
G302A
G303A
G304A
S305A
V306A
Q307A
I308A
V309A
Y310A

**Sup. Fig. 4. (LM)-CFP inclusions co-occur with (LM)-YFP aggregates.** Confocal microscopy of selected Tau RD (LM)-CFP variants 48 hours after lentiviral transduction of DS1 (A) and DS19 cells (B). Importantly, no inclusions were observed in DS1 cells which lack (LM)-YFP aggregates, while (LM)-CFP inclusions became apparent in cells with (LM)-YFP inclusions. Notably, not all variants contained (LM)-CFP inclusions (e.g. I308A and Y310A).

A

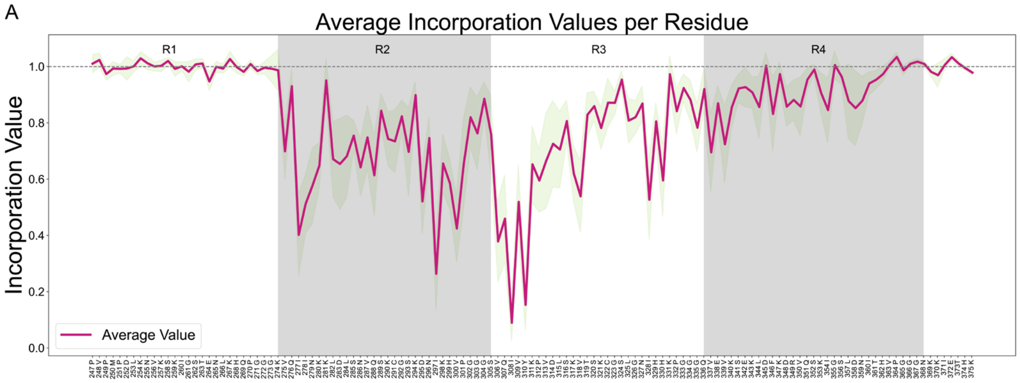
Average Incorporation Values per Residue
R1
R2
R3
R4
Incorporation Value
1.0
0.8
0.6
0.4
0.2
0.0
Average Value

**Sup. Fig. 5. Average Ala scan of all DS strains.** (A) Line plot of the average incorporation values for all strains revealed a distributed region that spans R2, R3 and R4 and was involved in aggregation in cells. Shaded area represents 95% confidence interval. Importantly, usage of the $^{306}$VQIVYK$^{311}$ region was a conserved feature among all strains. Throughout the length of the repeat domain, we found regions with patterns that involved every other residue, reminiscent of the expected alternating patterns in the beta-strands of an amyloid, for example: $^{293}$SDNIKHV$^{300}$, $^{305}$SVQIVYKPVDLSK$^{317}$and $^{326}$ GNIHHKP$^{332}$.

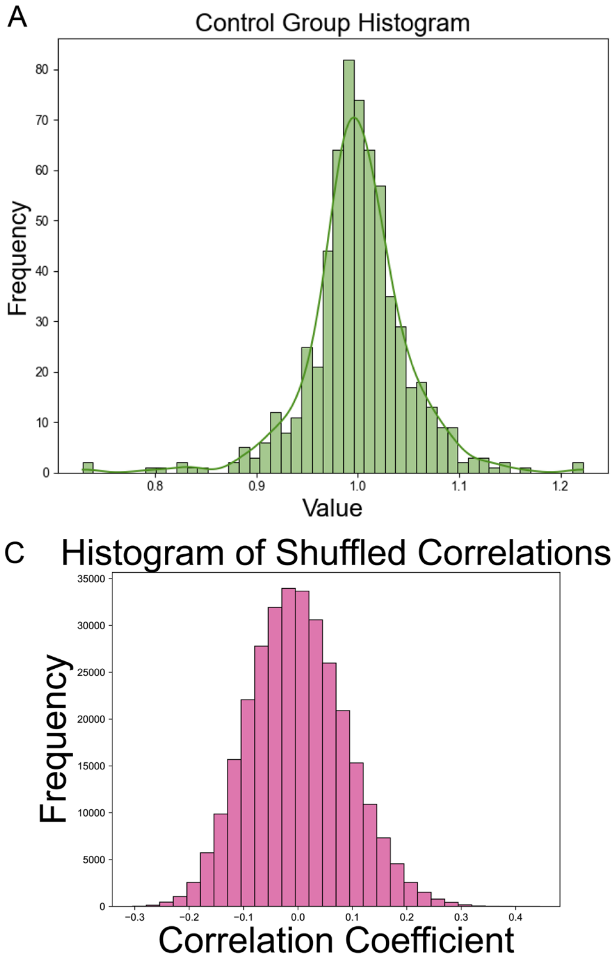

A
Control Group Histogram
Frequency
Value
B
Rest Group Histogram
Frequency
Value
C
Histogram of Shuffled Correlations
Frequency
Correlation Coefficient

**Sup. Fig. 6. Shuffled correlations for strain and residue.** To test the ability of the Ala scan to determine covariance of the scanned signature between strains, we considered the domains of tau RD that were not affected by Ala substitution (Control group: refers to non-variant residues, that is, the first 20 and last 10 amino acids of the sequence), and those that were (Rest Group: refers to the remaining residues). (A)The Control group values were randomly distributed around 1 in terms of their effect on incorporation. (B) The remaining sequences were highly skewed in their distribution, indicating that the population of hits were non-randomly distributed. (C) To test to what degree the observed correlations were meaningful, for each strain we generated 1000 permutations of independently shuffled values for each residue. This bounds the spurious correlations at an $R^2$ of 0.3, which is much lower than most of the observed correlations in Fig.4A.

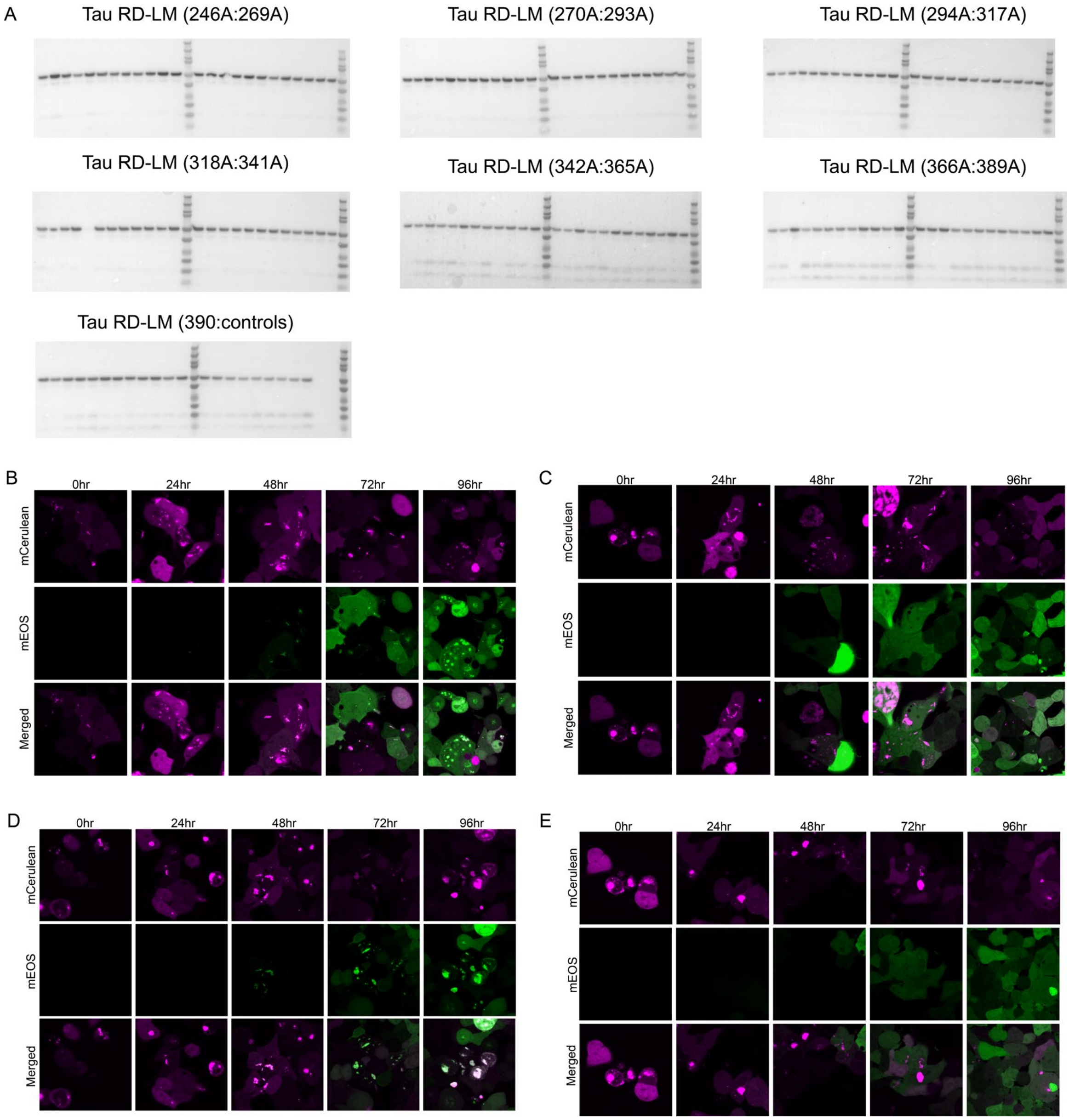
A
Tau RD-LM (246A:269A)
Tau RD-LM (270A:293A)
Tau RD-LM (294A:317A)
Tau RD-LM (318A:341A)
Tau RD-LM (342A:365A)
Tau RD-LM (366A:389A)
Tau RD-LM (390:controls)
B
C
D
E
0hr
24hr
48hr
72hr
96hr
mCerulean
mEOS
Merged

**Sup. Fig. 7. WT Ala scan validation.** (A) Western blots of 10μg of clarified cell lysate from HEK293T cells treated with all Tau RD(4Rext)-Cer alanine variants demonstrated robust expression of the fusion protein. Lanes were loaded with each variant in consecutive order. Tau protein was detected using an in-house rabbit polyclonal antibody, named Tau A, which targets R1. Confocal images of incorporation timecourse in: (B) AD seeded RD(3R/4Rext)-Cer/Ruby cells treated with WT-tau 246-408-mEOS3.2 (C) AD seeded RD(3R/4Rext)-Cer/Rub cells treated with 2P-tau 246-408-mEOS3.2. (D) AD seeded RD(4Rext)-Cer/Ruby cells treated with WT-tau 246-408-mEOS3.2. (E) AD seeded RD(4Rext)-Cer/Rub cells treated with 2P-tau 246-408-mEOS3.2. Importantly, the addition of the two proline mutations that inhibit tau's ability to aggregate prevented inclusion formation in cells. No puncta formed even after 96hrs of expression in cells that contained aggregates.

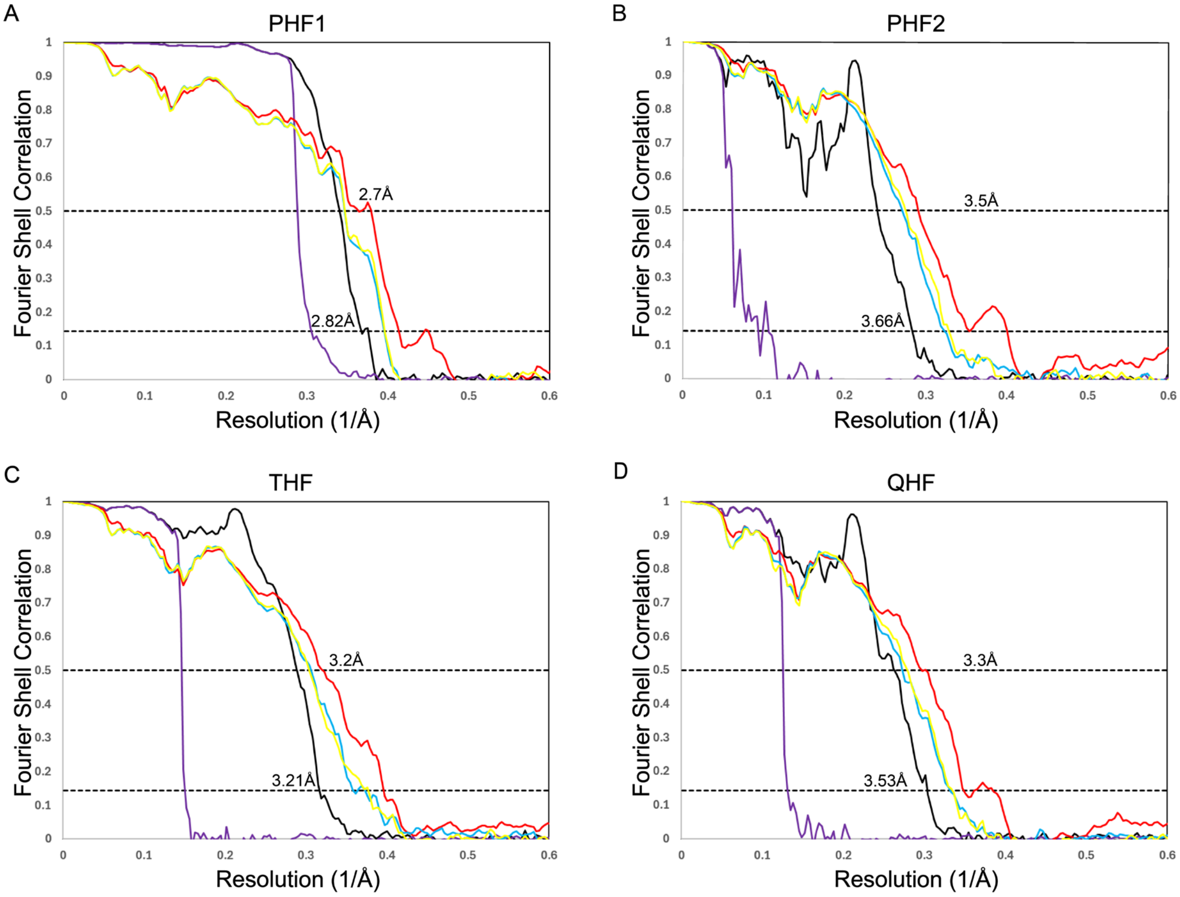


E

| Filament | Chain | RMSD to chain A of PDB:5o3L (Angstrom) |
|---|---|---|
| PHF1 | A | 0.897 |
| PHF1 | B | 0.902 |
| PHF2 | H | 1.32 |
| PHF2 | I | 1.376 |
| THF | L | 1.191 |
| THF | I | 2.541 |
| THF | F | 1.482 |
| QHF | A | 1.669 |
| QHF | B | 1.285 |
| QHF | K | 1.339 |
| QHF | L | 1.734 |

F

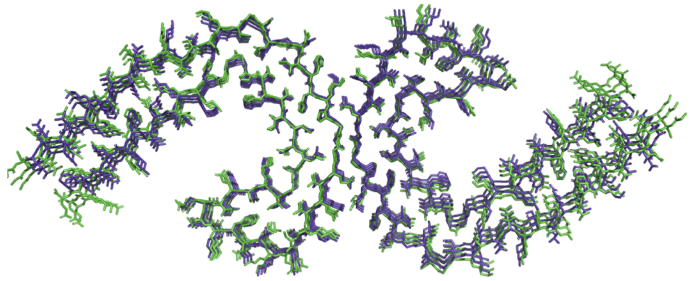

**Sup. Fig. 8. CryoEM reconstructions and atomic modeling.** Graphs illustrating the Fourier Shell Correlation (FSC) curves for the cryoEM models of tau (287-391) derived PHF1 (A), PHF2 (B), THF (C) and QHF (D), shown in black in each plot. Additionally, FSC curves are presented for the refined atomic model against the post-processed map utilized for atomic model generation (Red), for the refined atomic model from one half-map to itself (Blue), for the refined atomic model from the first half-map to the second (Yellow), and for phase randomized curves of the two independently refined half-maps (Purple). (E) Table showing the root mean-squared deviation of a single rung of each protofilament from the atomic models of tau (287-391) when compared to chain A of PDB 5o3l, which has the AD fold. Overlay of 3 rungs of PDB5o3l (purple) with 3 rungs of PHF1(green).

A

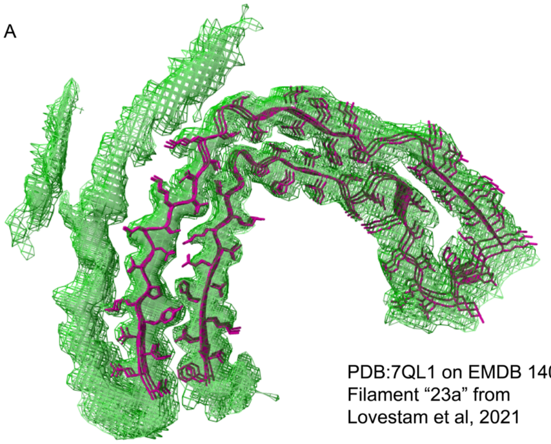

PDB:7QL1 on EMDB 14060
Filament “23a” from
Lovestam et al, 2021

B

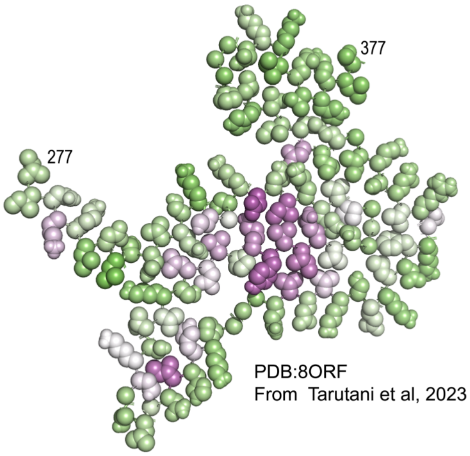


PDB:8ORF
From Tarutani et al, 2023

C

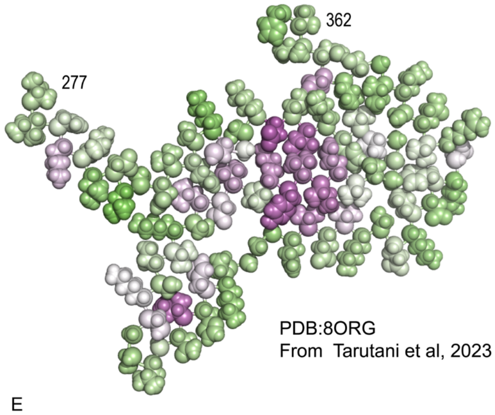


PDB:8ORG
From Tarutani et al, 2023

D

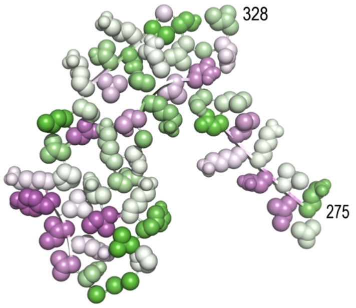


PDB:7QKW
Single protofilament
from Filament “44a” from
Lovestam et al, 2021

E

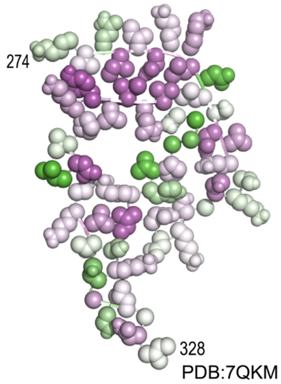


PDB:7QKM
Single protofilament
from Filament “45a” from
Lovestam et al, 2021

**Sup. Fig. 9. Positioning of Ala hits in relation to structures**. (A) *In vitro* assembled filaments made from tau (266-391) in PBS in the CTE fold, modeled in PDB:7QL1 into the density from EMDB 14060. The density threshold was set to highlight extra unmodeled density appearing to wrap back onto the core, presumably composed of the R2 sequence. (B and C). Two polymorphs reconstructed from filaments extracted from SH-S5Y5 cells expressing 1N4R tau and seeded with CBD brain homogenate. They were colored by the incorporation values measured for the average of all CBD cases. (D and E) PDB 7QKW of filament "44a" assembled from recombinant tau (266-391 S356D) in 10mM PB Ph7.4, 10mM DTT and 200mM KCl and PDB 7QKM of filament "45a" assembled from recombinant tau (266-391 S356D) in 10mM PB Ph7.4, 10mM DTT and 200mM NaCl. Both filament types were obtained in our preparation of Tau (266-391) in PBS with 10mM DTT (as in condition 23a from Lövestam et. al, 2022), importantly we did not observe filaments with the CTE fold and instead observed "44a" and "45a" in the absence of the S356D mutation. Both were colored by the incorporation values obtained from the preparation with these mixed filaments. Purple represents low incorporation and green represents no change in incorporation.

**Table S1. DS strain library**

| Strain | Inclusion Appearance | Origin |
|---|---|---|
| 2 | Mosaic | AGD cell line |
| 3 | Ordered | Recombinant Fibrils |
| 4 | Speckles | AD Homogenate |
| 5 | Speckles | Recombinant Fibrils |
| 6 | Threads | P301S Mouse Homogenate |
| 7 | Speckles | Recombinant Fibrils |
| 8 | Speckles | P301S Mouse Homogenate |
| 9 | Speckles | Recombinant Fibrils |
| 10 | Ordered | Recombinant Fibrils |
| 11 | Disordered | CBD Cell line |
| 12 | Speckles | CBD Cell line |
| 13 | Disordered | CBD Cell line |
| 14 | Ordered | Recombinant Fibrils |
| 15 | Threads | Recombinant Fibrils |
| 16 | Speckles | CBD Cell line |
| 17 | Speckles | AD cell line |
| 18 | Speckles | CTE Homogenate |
| 19 | Ordered | Recombinant Fibrils |

**Table S2. Subjects studied**

| Sample Name | ID | Age | Sex | Institution |
|---|---|---|---|---|
| AD1 | 49341 | 83 | F | UTSW |
| AD2 | 46879 | 76 | M | UTSW |
| AD3 | 46121 | 74 | F | UTSW |
| AD5 | 45408 | 73 | M | UTSW |
| 62579_Parietal | 62579 | 71 | M | WashU |
| 63377_Parietal | 63377 | 86 | M | WashU |
| 63870_Parietal | 63870 | 81 | F | WashU |
| 63870_Temporal | 63870 | 81 | F | WashU |
| CBD_12041 | 12044 | 73 | M | WashU |
| CBD1 | 48638 | 64 | M | UTSW |
| CBD2 | 44193 | 55 | M | UTSW |
| CBD5 | 49231 | 74 | M | UTSW |
| CBD6 | 46267 | 67 | F | UTSW |
| CBD7 | 37872 | 68 | M | UTSW |
| CBD_3950 | 3950 | 72 | M | WashU |
| CBD_23302 | 22302 | 62 | M | WashU |
| CTE_9130 | 9130 | | | Boston University |
| CTE_8408 | 8408 | | | Boston University |
| PSP2 | 45460 | 68 | F | UTSW |
| PSP1 | 47941 | 69 | M | UTSW |
| PSP3 | 49017 | 61 | F | UTSW |
| PSP4 | 48221 | 67 | M | UTSW |
| PSP_F1 | A06-231 | 76 | F | WashU |

### Table S3. CryoEM data collection and refinement statistics

| Cryo-EM data collection, refinement, and validation statistics | Truncated Tau 287-391 w/ 200mM MgCl2 | | | | Truncated Tau 297-391 w/ 200mM MgCl2 | | | Truncated Tau 266-391 w/ 200mM NaCl | |
|---|---|---|---|---|---|---|---|---|---|
| Electron Microscope Type | Titan Krios | | | | Titan Krios | | | Titan Krios | |
| Nominal Magnification | 105,000x | | | | 105,000x | | | 105,000x | |
| Voltage (kV) | 300 | | | | 300 | | | 300 | |
| Detector | Gatan K3 w/ Bioquantum Energy Filter | | | | Gatan K3 w/ Bioquantum Energy Filter | | | Gatan K3 w/ Bioquantum Energy Filter | |
| Electron Exposure (e-/Å²) | 57 | | | | 52 | | | 62 | |
| Defocus Range (μm) | -0.8 to -2.2 | | | | -1.0 to -2.2 | | | -1.5 to -3.0 | |
| Super-resolution Pixel Size (Å) | 0.415 | | | | 0.415 | | | 0.415 | |
| **Reconstruction** | **PHF** | **THF** | **QHF** | **PHF2** | **PHF** | **THF** | **QHF** | **44a** | **45a** |
| Total number of micrographs | | | | | | | | | |
| Number of usable micrographs | 2,277 | | | | 5,094 | | | 2,528 | |
| Particles after Extraction | 336,719 | | | | 288,528 | | | 348,124 | |
| Particles after 2D Classification | 163,219 | | | | | | | 142,881 | 137,756 |
| Particles after 3D Classification | 152,026 | 41,201 | 20,671 | 16,801 | 50,023 | 28,122 | 80,078 | | |
| Map Resolution (Å, FSC=0.143) | 2.82 | 3.21 | 3.53 | 3.66 | 3.21 | 3.32 | 2.79 | 4.41 | 3.86 |
| Helical Rise (Å) | 2.37 | 4.75 | 2.37 | 4.75 | 2.38 | 4.75 | -0.81 | 4.82 | 2.41 |
| Helical Twist (°) | 179.47 | -0.86 | 179.58 | -0.87 | 179.5 | -0.86 | 4.75 | -0.93 | 179.58 |
| Symmetry Imposed | 2-1 | C1 | 2-1 | C1 | 2-1 | C1 | C1 | C2 | C1 |
| EMDB Entry | EMD-46908 | EMD-46911 | EMD-46915 | EMD-46909 | EMD-46870 | EMD-46885 | EMD-46904 | EMD-47092 | EMD-46937 |
| **Refinement** | | | | | | | | | |
| Initial model used (PDB Code) | 5o3l | 5o3l | 5o3l | 5o3l | | | | 7QKW | 7QKM |
| Model resolution (Å, FSC=0.5) | 2.7 | 3.2 | 3.3 | 3.5 | | | | | |
| Map sharpening B factor (Å²) | -52.32 | -62.94 | -59.15 | -65.17 | | | | | |
| Model composition | | | | | | | | | |
| Non-hydrogen atoms | 4,419 | 6,625 | 14,371 | 4,416 | | | | | |
| Protein residues | 462 | 693 | 924 | 462 | | | | | |
| Ligands | N/A | N/A | N/A | N/A | | | | | |
| B Factors (Å²) | | | | | | | | | |
| Protein | 72.95 | 72.59 | 72.95 | 72.95 | | | | | |
| Ligand | N/A | N/A | N/A | N/A | | | | | |
| R.m.s. deviations | | | | | | | | | |
| Bond lengths (Å) | 0.010 (0) | 0.011 (0) | 0.011 (0) | 0.011 (0) | | | | | |
| Bond angles (°) | 1.941 (20) | 2.024 (40) | 1.982 (37) | 2.056 (26) | | | | | |
| Validation | | | | | | | | | |
| MolProbity score | 0.95 | 1.12 | 1.15 | 1.41 | | | | | |
| Clashscore | 0 | 0 | 0.07 | 0 | | | | | |
| Poor rotamers (%) | 0 | 1.16 | 0.75 | 1.74 | | | | | |
| Ramachandran plot | | | | | | | | | |
| Favored (%) | 92.89 | 89.33 | 86.78 | 80.44 | | | | | |
| Allowed (%) | 7.11 | 10.67 | 13.22 | 19.56 | | | | | |
| Disallowed (%) | 0 | 0 | 0 | 0 | | | | | |

**Other Supplementary Materials for this manuscript include the following:**

Movies S1: **cryo-ET of DS13-extracted filaments**

Data S1: **Source data.**
